# UVB-Induced Genotoxic Stress Activates the DNA Damage Response and Innate Immune Pathways in Sea Urchin Coelomocytes

**DOI:** 10.64898/2026.01.14.699502

**Authors:** Riss M. Kell, Margaret K. Weber, Rosa Y. Escalante, Jeffrey C. Silva, Andrea G. Bodnar

## Abstract

Mechanistic crosstalk between the DNA-damage response (DDR) and the innate immune system is essential to maintain genomic integrity, tissue homeostasis, and organismal resilience. However, the origin and diversity of this crosstalk among animal lineages remain poorly understood. Here, we used the purple sea urchin *Strongylocentrotus purpuratus* to identify immune-cell-intrinsic responses to genotoxic stress. This species is a long-lived, cancer-resistant, basal deuterostome that diverged from the vertebrate lineage ∼550 million years ago, providing a powerful system to investigate the ancestral origins and complexity of DDR-immune system crosstalk. UVB-induced DNA damage elicited a robust transcriptional response in *S. purpuratus* immune cells (coelomocytes), activating conserved DDR genes in addition to autophagy, ubiquitin signaling, and innate immune pathways. Single-cell RNA sequencing at six hours post UVB exposure identified phagocytes and vibratile cells as the major mediators of the response to UVB challenge. Functional assays and western blots confirmed increased autophagy and widespread post-translational modification (ubiquitination and phosphorylation) of substrates in key signaling pathways. Our findings demonstrate the concurrent induction of DDR and innate immune pathways in an invertebrate deuterostome and establish a molecular framework for understanding how echinoderms respond to damaged self in the absence of vertebrate-specific DNA sensors. By revealing the conserved mechanisms that link the DDR with innate immunity, this work lays the foundation for comparative studies of damage recognition systems that shape deuterostome defenses, promoting genomic stability and cancer resistance.

## 1 Introduction

Cells are continuously challenged with DNA damage induced by both exogenous (*i.e.,* radiation, carcinogens) and endogenous (*i.e.,* replication errors, reactive oxygen species) insults (1–3). DNA lesions must be addressed promptly to maintain cellular homeostasis and avoid genomic instability and oncogenesis. To preserve genomic integrity, cells are equipped with a complex network referred to as the DNA-damage response (DDR) to detect DNA lesions and coordinate repair, senescence, or apoptosis (4,5). It has become increasingly recognized that the DDR and the immune system engage in important crosstalk to maintain genomic fitness (6–8). The immune system has core functions in the recognition and clearance of damaged cells by detecting damage-associated molecular patterns (DAMPs) through pattern recognition receptors (PRRs) (Tang et al. 2012; Nastasi et al. 2020). Key DAMP-sensing PRRs include Toll-like receptors (TLRs), NOD-like receptors (NLRs), and AIM2-like receptors (ALRs), which trigger downstream signaling to orchestrate immune responses (9).

Cytosolic or endosomal self-DNA acts as a DAMP that is recognized by DNA sensors that induce type I interferons (IFNs) or inflammasomes (10). The cGAS-STING pathway has emerged as one of the key DNA sensors in vertebrates, driving IFN responses upon detecting cytosolic DNA (11,12). Other cytosolic DNA sensors include AIM2-like receptors and Z-DNA-binding protein 1 (ZBP1), which activate inflammasome pathways that trigger pyroptosis, a lytic form of cell death (10,13). Furthermore, DEAH- and DEAD-box helicases such as DHX9, DHX36, and DDX41, as well as RNA polymerase III contribute to DNA-mediated innate immunity, while endosomal DNA is recognized by TLR9, which induces IFNs and inflammatory cytokines through immune response factor 7 (IRF7)–nuclear factor-κB signaling (NF-κB) signaling (10,13). Although progress has been made in understanding the coordination between the DDR and immunity in mammals, especially regarding the cGAS-STING axis, the origins and diversity of DDR-immune crosstalk among animal lineages remain poorly understood.

Echinoderms have played a foundational role in the history of immunology, providing early insights into self/non-self discrimination and revealing unexpected complexity within the echinoderm innate immune system (14,15). Building on this legacy, echinoderms such as sea urchins offer a powerful model for discovering novel interactions between the DDR and the innate immune system. Sea urchins are a well-established animal model with a long history of contributions to understanding fundamental biological processes including cell cycle regulation and the gene regulatory networks of early development (16–18). Despite a wealth of data on their life histories and incidence of disease (19,20), and in contrast with numerous reports of neoplasia in other commercially fished marine invertebrates such as oysters, mussels, and clams (21), sea urchins are noted for their absence of neoplastic disease (22–27). The lack of observed neoplasia is particularly striking given that sea urchins possess significant regenerative capacity and that some species are exceptionally long-lived with lifespans in excess of 100 years (23). This suggests that sea urchins possess robust mechanisms to maintain genomic integrity and prevent tumorigenesis. As basal deuterostomes, sea urchins lack V(D)J-based, Rag1/2-mediated adaptive immunity, yet possess a remarkably complex innate immune system that includes expanded families of PRRs (including TLRs, NLRs, and RLRs), conserved effector pathways involving NF-κB, IKKε, TBK1, IL-17, and TNF, and diverse anti-microbial peptides (15,28–31). Notably, sea urchins lack canonical components of the vertebrate DNA sensing pathway cGAS-STING (i.e. cGAS, STING, IRF3, and IRF7) (32,33), suggesting that these animals have alternative or ancestral immune strategies for recognizing and responding to endogenous DNA damage.

Sea urchin immune effector cells, collectively referred to as coelomocytes, are classified into four major cell types: phagocytes, red and colorless spherule cells (RSCs and CSCs; also referred to as red and colorless amoebocytes), and vibratile cells (14,15). Coelomocyte subpopulations are capable of phagocytosis, chemotaxis, and cytotoxic responses to injury or infection (34–37), functions that are analogous to those of vertebrate granulocytes, macrophages, and natural killer cells. The phagocyte class of sea urchin coelomocytes is morphologically and functionally complex. These cells can be subclassified into large, medium, and small phagocytes based on size, and large phagocytes can be further resolved into polygonal or discoidal types based on cytoskeletal morphology (38–41). Each phagocyte subtype exhibits distinct levels of substrate binding and phagocytic activity (42). Red spherule cells are so named due to their cytoplasmic vesicles containing the naphthoquinone pigment echinochrome A (EchA), which possesses antimicrobial activity and functions in innate defense by chelating iron (43). Colorless spherule cells and vibratile cells are comparatively understudied, and the response of any individual coelomocyte cell type to genotoxic challenge has not been characterized.

Sea urchins possess a highly conserved DDR, which involves the detection of DNA damage, activation of cell cycle checkpoints, engagement of DNA repair pathways, or induction of apoptosis in response to genotoxicant exposure (44–46). A link between the innate immune system and the DDR has been previously suggested by the observation that genotoxic stress induces the expression of innate immune genes in *Lytechinus variegatus* coelomocytes (47). Long-lived sea urchin species such as the purple sea urchin (*S. purpuratus*, lifespan > 50 years) (48–50), and the red sea urchin (*Mesocentrotus franciscanus*, lifespan > 100 years) (23), possess expanded repertoires of innate immune genes compared to shorter-lived species (30,33), suggesting augmented functions in innate immunity to support long-term genomic stability and cancer resistance. Yet, despite the conserved DDR and complex innate immune system in sea urchins, the molecular mechanisms by which sea urchin coelomocytes respond to DNA damage remain largely unexplored. In particular, the contributions of distinct coelomocyte cell types to the DDR, and how this response intersects with innate immune signaling, are unknown. Here, we exposed coelomocytes of the long-lived purple sea urchin (*S. purpuratus*) to UVB radiation *in vitro* and characterized the response over a timecourse of recovery using bulk RNA sequencing, single-cell RNA sequencing, and functional assays. This approach provided a cell-type-resolved view of the sea urchin DDR, revealing the intrinsic capacities of immune effector cells to detect and respond to genotoxic stress. Our findings underpin a framework for understanding how these mechanisms may contribute to genomic stability and cancer resistance in this long-lived sea urchin species.

## 2 Materials and Methods

### 2.1 Animal husbandry

*Strongylocentrotus purpuratus* were purchased from a commercial vendor (Marinus Scientific) and housed in recirculating aquaria containing artificial seawater (Instant Ocean, 33 ppt salinity) set at ambient temperature (13°C). Upon receipt, animals were acclimated to these conditions for 1 month to recover from the stress of initial collection and to ensure that all animals were subjected to identical environmental conditions prior to experimentation. Animals were maintained on a diet of rehydrated dried kelp (*Laminaria digitata*) purchased from a commercial vendor.

### 2.2 Challenge of human PBMCs with UVB

Human peripheral blood mononuclear cells (PBMCs, ATCC #PCS800-011) were purchased from the American Type Culture Collection (ATCC, Manassas, VA). Immediately prior to use, the cell stock vial was removed from liquid nitrogen storage, quickly thawed in a 37°C water bath until only a small amount of ice remained, and transferred to a 15 mL sterile Falcon tube. Cells were gently resuspended with 10 mL of thawing media (Hank’s Balanced Salt Solution without Ca^2+^ or Mg^2+^, ATCC #30-2213) supplemented with 10% fetal bovine serum (FBS). A sample of cells was taken for an initial count and viability assessment using the trypan blue exclusion assay. Cells were diluted at 1:1 volume in 0.4% (w/v) trypan blue in Dulbecco’s phosphate buffered saline (DPBS) and manually counted using a hemocytometer. Cells were pelleted by centrifugation (300 x g, 5 min, 4°C) and the supernatant was removed. Cells were resuspended in RPMI 1640 media (ATCC #30-2001) supplemented with 10% FBS, transferred into a T75 flask at 2.1 x 10^6^ cells/mL, and incubated at 37°C with 5% CO_2_ for 24 h to allow cells to recover from cryopreservation. Cells were refreshed with new RPMI 1640 containing 10% FBS before 200 µL (1.76 x 10^5^ cells) was aliquoted into sterile 1.5 mL Eppendorf tubes. Cells were irradiated using 0, 1000, 2000, 4000, 6000, and 9999 mJ/cm^2^ UVB radiation (n = 3 technical replicates per dose) using an Analytik Jena UVP Crosslinker CL-3000 to induce DNA damage before transferring to a sterile 24-well tissue culture plate. The plate was covered with foil to prevent incident light damage and incubated at 37°C with 5% CO_2_. Cell viability was assessed by trypan blue exclusion assay after 24 h. Cells were diluted at 1:1 volume in 0.4% (w/v) trypan blue in DPBS. Viability measurements were used to calculate LD_50_ values at 24 h recovery using the AAT Bioquest LD_50_ calculator in three-parameter mode (https://www.aatbio.com/tools/ld50-calculator) (51).

### 2.3 Coelomocyte collection and challenge with UVB

A sterile 21-gauge needle fitted to a sterile 5 mL syringe barrel was used to pierce through the peristomial membrane of *S. purpuratus* (n = 3) into the coelomic cavity and withdraw 1 mL of whole coelomic fluid (WCF). To avoid cell shear, the needle was removed from the syringe barrel prior to gently expelling the WCF into sterile 1.5mL Eppendorf tubes. Samples were immediately placed on ice and covered with foil to prevent incident light damage. Samples were irradiated at 0, 1000, 2000, 4000, 6000, and 9999 mJ/cm^2^ UVB radiation using an Analytik Jena UVP Crosslinker CL-3000. Cells were allowed to recover for 24 h at 17°C in Eppendorf tubes, covered with foil to prevent incident light damage. Cell viability was assessed using the trypan blue exclusion assay. Cells were gently resuspended and diluted at 1:1 volume into anticoagulant (Ca^2+^ and Mg^2+^-free seawater containing 30 mM EDTA: 460 mM NaCl, 10 mM KCl, 7 mM Na_2_SO_4_, 2.4 mM NaHCO_3_, pH 7.4; CMFSW-E). An aliquot of anticoagulated cells was further diluted 1:1 (v/v) into 0.4% (w/v) trypan blue in CMFSW-E before loading and counting on a hemocytometer.

### 2.4 Quantification of DNA damage using the comet assay

DNA damage in *S. purpuratus* coelomocytes resulting from 1,000 mJ/cm^2^ UVB was assayed using the OxiSelect Comet Assay kit (Cell Biolabs Inc, #STA-351). All reagents were prepared according to the manufacturer’s protocol. WCF was collected, UVB treated, and allowed to recover *in vitro* as described above. WCF (50 μL) was diluted 1:1 (v/v) in anticoagulant (CMFSW-E), and samples were centrifuged at 700 x g for 2 min before the supernatant was discarded. The cell pellet was resuspended in 500 µL CMFSW before centrifuging at 700 x g for 2 min and discarding the supernatant. Cells were resuspended in 500 µL of CMFSW. Cell concentration and viability were evaluated using the trypan blue exclusion technique as detailed above. Cells were resuspended in CMFSW-E to a final concentration of 1 x 10^5^ cells/mL. Seven µL of this cell suspension was mixed with 70 µL of 37°C 1% low melting point agarose (LMPA) before adding onto a 3-well comet slide (Cell Biolabs). Each slide held three biological replicates. Slides were coverslipped (VWR #48393-059) and placed on ice for 15 min to solidify gels before coverslips were removed. Cells were lysed by placing slides into 4°C lysis buffer (10 mM Tris, 100 mM EDTA, 2.5 M NaCl, 10% DMSO, 10% Triton X-100, pH 10) for 1 h on ice. Slides were submerged in 4°C alkaline buffer (0.3 M NaOH, 1 mM EDTA, pH > 13) for 40 min to denature the DNA. Slides were electrophoresed in alkaline electrophoresis buffer (0.3 M NaOH, 1 mM EDTA) at 20 V, 300 mA, for 25 min. Ice was packed around the electrophoresis chamber to keep the buffer cold. Slides were washed using three 5 min Milli-Q water immersions. Slides were incubated in 70% ethanol for 5 min before air drying for 1 h at room temperature and staining with VistaGreen DNA dye (Cell Biolabs) for 15 min. Comet images were taken at a magnification of 4x at maximum resolution (2048 x 2048 pixels) using a fluorescence microscope (Olympus IX83) fitted with an X-CITE 120LED Boost System and FITC filter cube (excitation filter of 475/50 nm, emission filter of 540/50 nm). Microscopy images were analyzed using AutoComet (Barbé et al. 2023). A minimum of 50 cells from each of three biological triplicates was scored. Average percentage of DNA in the comet tail relative to the nuclear region was used to quantify DNA damage.

### 2.5 Bulk RNA extraction and sequencing

WCF was collected from *S. purpuratus* (n = 3) and treated samples were dosed with 1,000 mJ/cm^2^ UVB radiation as detailed in section 2.3 and allowed to recover *in vitro* at 17°C. At 0, 1, 3, 6, and 24 h recovery, a 500 µL aliquot of WCF (∼5 x 10^5^ cells) was collected for bulk RNA-seq analysis. Aliquots were centrifuged at 5,000 x g for 5 min at 13°C and the supernatant was discarded. Cell pellets were stored at -80°C until further analysis. Total RNA from 30 cell pellets (3 sea urchins x 5 timepoints x 2 conditions) was extracted using the RNeasy Plus Mini Kit (Qiagen, #74134) following the manufacturer’s instructions with the following modifications: i) 200 µL of buffer RLT+ containing beta-mercaptoethanol was added to each pellet, ii) pellets were mechanically homogenized using a diethyl pyrocarbonate (DEPC)-treated (RNAse-free) sterile plastic pestle, and iii) 400 µL of additional buffer RLT+ containing beta-mercaptoethanol was added before vortexing the sample for 30 sec. Extracted RNA was eluted in 30 µL of RNAse-free water. Eluted RNA concentrations were determined using a Qubit 4 Fluorometer and Qubit RNA high-sensitivity assay (Thermo Fisher Scientific, #Q32852). RNA quality in each sample was assessed using a 5300 Fragment Analyzer System (Advanced Analytical) and HS RNA (15 nt) kit (Agilent, #5191-6574) using 2000 pg input RNA following the manufacturer’s instructions. Samples were sent to the Center for Genome Innovation at the University of Connecticut (UCONN-CGI; Storrs, CT) for library preparation using a Low Input mRNA SMART-Seq LP kit (Takara Bio). Samples were sequenced (50M total paired end reads per sample, 100 bp reads) on an Illumina NovaSeq 6000 with a NovaSeq S4 200 cycle sequencing kit (Illumina). Sample quality was assessed using FastQC v0.12.1 (52).

### 2.6 Bulk RNA data analysis

Sequences were trimmed with Trimmomatic v0.39 (53) (SLIDINGWINDOW: 4:15, MINLEN: 50). The trimmed reads were mapped to the *S. purpuratus* genome (v5.0) using HISAT2 (v2.2.1) (54). Both properly paired reads and unpaired forward reads were retained for downstream analysis. Of 30 total samples, one sample was removed from further analysis (UVB-treated, 3 h recovered, replicate A) based on poor percentage of surviving reads (55%) compared to all other samples (> 75% survival). Gene count files were created using HTSeq v2.0.4 (55) using the HISAT2 alignment files and the *S. purpuratus* v5.0 Gene Transfer Format (GTF) file. HTSeq count files were combined into a single table in R and genes with zero counts across all samples were removed. Further data processing was performed based on Chille et al. 2022 (56). The gene count matrix was filtered to remove low coverage counts using the pOverA function of the ‘genefilter’ package v1.86.0 (57). For a gene to pass pOverA filtering, it required a minimum count of 10 in at least three samples of 29 total samples (pOverA = 3/29 = ∼0.1; expression threshold = 10). Differential gene expression was performed in R using DESeq2 v.1.44.0 (58) comparing biological triplicate control and UVB-treated samples at each individual timepoint. Results were filtered to retain only significantly differentially expressed genes (DEGs) (Benjamini-Hochberg adjusted *p* values < 0.05, log_2_ fold-change > 1 or < - 1). This resulted in 2,581 unique DEGs inclusive of all timepoints. Gene matrices were transformed with the DESeq2 variance stabilizing transformation (VST) function after confirming that all size factors were less than four. Genes were grouped into two clusters using k-means clustering in R. To visualize the bulk RNA sequencing data in a heatmap, the filtered, VST-transformed gene count matrix was first centered by subtracting the mean expression value of each row from each element in that row. Data was visualized using the ‘Heatmap’ function from the ComplexHeatmap package (v.2.20.0) (59). Gene count plots showing trends in mean VST gene expression counts for Cluster 1 and Cluster 2 genes were generated using group means with error bars representing ±1 standard error of the mean (±1 SEM) calculated from n = 3 biological replicates per treatment (with the exception of the UVB-treated, 3 h recovered group, which had n = 2 biological replicates as mentioned above) per time point. A two-way ANOVA in R was used to determine whether there were significant main effects of treatment, time, or their interaction (significance was set to *p* < 0.05).

### 2.7 Bulk RNAseq: pathway enrichment analysis using STRING

A protein FASTA of Cluster 1 and Cluster 2 protein-coding genes was created by matching gene IDs to the *S. purpuratus* protein FASTA sequences available through EnsemblGenomes ( https://ftp.ensemblgenomes.ebi.ac.uk/pub/metazoa/release-60/fasta/strongylocentrotus_purpuratus/pep, last modified 2024-08-09, last accessed 2025-03-24).

Protein FASTA sequences were used as input for enrichment analysis within STRING (“Proteins by sequences” input; https://string-db.org/) using *S. purpuratus* as the reference organism (60). STRING parameters employed the highest confidence settings (0.900 minimum required interaction score) with all interaction sources selected (text mining, experiments, databases, co-expression, neighborhood, gene fusion and co-occurrence). Enriched terms are listed in Supplementary Table 1.

### 2.8 Single-cell RNA sequencing

WCF was collected, and cells were challenged with 1,000 mJ/cm^2^ UVB and allowed to recover *in vitro* as detailed above. At 6 h recovery, a 500 µL aliquot (∼5 x 10^5^ cells) was collected for single-cell RNA-seq (scRNAseq). Samples were processed for scRNAseq using the 10x Genomics Chromium Controller platform and 10x Genomics Chromium NextGEM Single Cell v3.1 protocol

(61). Samples were decoagulated by adding ice-cold CMFSW-E (pH adjusted to 8.2) to cells at 1:1 volume with gentle pipetting. Samples were centrifuged at 400 x g for 3 min at 13°C and the supernatant was discarded. Cell pellets were gently resuspended in 500 µL cold hypertonic PBS (1X PBS with an additional 350 mM NaCl) as reported previously in the preparation of cells from marine animals for scRNAseq (62). Samples were centrifuged at 400 x g for 3 min at 13°C and the supernatant was discarded. Cell pellets were resuspended in 500 µL of hypertonic PBS. Cell density and viability were assessed via the trypan blue exclusion method. The resuspended cell solution (10 µL) was combined with 10 µL of 0.4% trypan blue (w/v in hypertonic PBS) prior to manually counting cells using a hemocytometer. Samples were further diluted in hypertonic PBS to appropriate loading concentrations according to Chromium Next GEM Single Cell 3’ v3.1 instructions targeting 10,000 cells per sample. Eight samples (four controls, four UVB treated) were loaded into individual lanes of a Chromium NextGEM Chip G. cDNA amplification and library prep were performed according to the manufacturer’s protocols. The average cDNA library fragment size for each sample (∼400 nt) was determined using a 5300 Fragment Analyzer System (Advanced Analytical) and HS NGS Fragment (1-6000 bp) kit (Agilent, #5191-6578) following the manufacturer’s instructions. Final cDNA libraries were sequenced at UCONN CGI. Sample libraries were sequenced to 30,000 read pairs per cell (60,000 total reads) on an Illumina NovaSeq 6000 with Illumina NovaSeq SP 100 cycle v1.5 and NovaSeq S1 100 cycle v1.5 sequencing kits. The quality of the resulting FASTQ files was confirmed using FastQC v0.12.0 (52). FASTQ files were uploaded to the 10x Genomics Cloud Analysis platform (https://cloud.10xgenomics.com/cloud-analysis) using the 10x Genomics Cloud CLI for Linux (https://www.10xgenomics.com/support/software/cloud-analysis/latest/tutorials/CA-cloud-cli-documentation-for-linux). A custom reference genome was created using CellRanger v7.1.0 ‘mkref’ using the *S. purpuratus* genome (v5.0) genome FASTA and GTF annotation file as inputs. The GTF file was filtered to contain only protein-coding genes using the CellRanger ‘mkref’ function. The CellRanger ‘count’ function (v7.1.0) was used to align sequencing reads to the *S. purpuratus* reference genome (v5.0) within the 10x Genomics cloud analysis platform to create a raw feature/cell matrix HD file for each sample. Two samples (one control and its corresponding UVB-treated sample) were excluded from further analysis due to a wetting failure in the 10x hardware during GEM generation.

### 2.9 Single-cell RNAseq data analysis

#### 2.9.1 Preprocessing

Ambient RNA, doublets, and low-quality sequence data from individual cells were removed during data preprocessing. Ambient RNA were identified and removed from all raw data files (HDF5 files) using the CellBender (v0.3.0) ‘remove-background’ algorithm (63) with the following parameters: 50,000 total droplets included, 10,000 expected cells, 0.01 fpr, 150 epochs. Output .h5 files were used for further analysis in Python 3.11 using Jupyter Notebook v7.1.3 (64), Scanpy v1.10.1 (Wolf et al. 2018), and SCVI tools v1.1.2 (65). Doublets were identified using DoubletDetection v2.4 (https://zenodo.org/records/14827937) and removed from each scRNA-seq count matrix. Cells with total feature counts falling beyond five absolute deviations of the median were removed as outliers (66).

#### 2.9.2 Integration, clustering, and normalization

Processed .h5 files were combined into one adata object using the Scanpy ‘concat’ function. The Scanpy ‘pp.filter_genes’ function was used to remove genes found in less than 50 cells. The adata object was passed to the SCVI ‘model.train’ function with default parameters to perform data integration (107 epochs). The model was saved to a .model file. The SCVI ‘model.get_latent_representation’ function was used to acquire embeddings, and neighbors were calculated using the Scanpy ‘sc.pp.neighbors’ function. Leiden clustering was performed using the Scanpy ‘sc.tl.leiden’ function at a resolution of 1. A new adata layer called ‘counts’ was saved by creating a copy of adata.X (the raw, unnormalized data) prior to normalization, such that raw data could still be accessed for downstream differential expression analysis. Data normalization was performed using the Scanpy ‘pp.normalize_total’ function prior to log transformation using the ‘pp.log1p’ function (67). A new adata layer called ‘condition’ was created in adata.obs such that every cell was labelled by sample condition. Clusters were colored by condition, leiden cluster, and assigned cell type using the Scanpy ‘pl.umap’ function.

#### 2.9.3 Cell type assignment using marker genes

A cell type dictionary was prepared relating each Leiden cluster number to one of four cell types (phagocytes, vibratile cells, red spherule cells, and colorless spherule cells). Differential expression was performed to identify those genes that were differentially expressed (adjusted *p* value < 0.05, log fold-change > 0.5) among all clusters using the Scanpy functions ‘tl.rank_genes_groups(adata, groupby = ‘leiden’)’ and ‘get.rank_genes_groups_df’. The differentially expressed genes were saved to a dataframe called ‘markers’. To identify clusters as specific cell types, the markers dataframe was searched for significant (adjusted *p* value < 0.05) expression of the following marker genes: *Pks1* for red spherule cells; *SpTrf, Sp-B7L3*, toll-like receptors (TLRs), and complement C3 for phagocytes; *Sp-P2rx4*, *FoxJ1*, and dynein heavy chain genes for vibratile cells; and *DD104*, lysozyme, and strongylocin for colorless spherule cells (Supplementary Table 4). The total number of cells in each sample was quantified, and cells were grouped by sample, condition, and cell type to calculate cell counts. Frequencies of each cell type within conditions were determined by dividing the cell-type-specific counts by the total number of cells per sample. Cell type frequencies were visualized using boxplots and swarmplots in Python (matplotlib and seaborn).

#### 2.9.4 Differential expression analysis

Differential expression (DE) analysis was performed using the SCVI tools ‘model.differential_expression’ function on the raw count matrix stored in adata.X. The output dataframe was filtered to include only those genes with a false discovery rate < 0.05. The following columns were added to the output dataframe to facilitate plotting: a column of log_10_ *p* values, a column of negative log_10_ *p* values, and columns for descriptive gene name, Gene ID, NCBI taxon ID, Entrez ID, Uniprot ID, and Ensemble IDs that were mapped to LOC IDs using file GeneExternalRef.txt downloaded from Xenbase (https://download.xenbase.org/echinobase/GenePageReports/GeneExternalRef.txt). Bokeh volcano plots were created to visualize differentially expressed genes. Significantly upregulated genes were those with a negative log_10_ (*p* value) > 1.3 and a log_2_ fold-change mean > 1. Significantly downregulated genes were those with a negative log_10_ (*p* value) > 1.3 and a log_2_ fold-change < -1.

#### 2.9.5 STRING analysis

A protein FASTA for each list of differentially expressed, upregulated genes (UDEGs) was created by mapping UDEG LOC gene identifiers to LOC identifiers in the *S. purpuratus* v5.0 peptide FASTA (https://www.echinobase.org/echinobase/static-echinobase/ftpDatafiles.jsp). GO, KEGG, and Reactome enrichment analysis and protein-protein interactions among the UDEGs specific to each cell type were investigated using STRING v12.0 (68) using the high confidence setting (0.700 minimum required interaction score) with all interaction sources selected (text mining, experiments, databases, co-expression, neighborhood, gene fusion and co-occurrence) and *S. purpuratus* as the target organism.

#### 2.9.6 Quantification of autophagy

WCF was collected from n = 3 *S. purpuratus*, expelled into low binding microfuge tubes (Eppendorf, #EP02243108), and cell counts and viability assessments were performed as detailed in section 2.4. Cells were treated with 1000 mJ/cm^2^ UVB using an Analytik Jena UVP Crosslinker (CL-3000M) and allowed to recover in vitro at 17°C for 6 h. The negative control cells were collected and treated similarly but the exposure to UVB was omitted. Detection and quantification of the autophagic signal was performed using an Autophagy Assay Kit (Abcam, ab139484) following the manufacturer’s instructions with the following modifications. The anticoagulant was supplemented with 5% FBS and used in place of 1x Assay Buffer. Cells were stained for 30 min at 17 with a combination of green detection reagent (1:1000) and Hoechst 33342 (1:1000) that targeted autophagic vesicle receptors and nuclei, respectively. After staining, 7.5 x 10^4^ cells per sample were seeded into a clear cell culture treated 96-well plate (Costar, #3599). Live cell imaging (40X magnification) was performed with an Olympus IX83 inverted fluorescent microscope using DAPI and FITC filters. Fluorescence was quantified using ImageJ (1.54p) with Java 1.8.0 (69). Threshold values were automatically generated for each FITC image such that fluorescing probes were included in the region of interest (ROI) while the background was excluded using triangle thresholding (70). Area-averaged integrated fluorescence density was then measured. Nuclei were counted for each Hoechst 33342 image by Otsu thresholding (71) and watershed separation to count individual particles. FITC integrated density values were normalized using nuclei (Hoechst 33342) counts for each image. FITC-normalized integrated density values were subject to a Shapiro-Wilk test of normality (*p* = 0.007, indicating data was not normally distributed, therefore a non-parametric test was used for statistical testing). A Wilcoxon rank sum test was used to test for significant differences between UVB-treated groups and the controls (biological triplicate samples, analyzed in technical triplicate for n = 9 images analyzed per treatment).

#### 2.9.7 Western blot

WCF (10 mL) was withdrawn from n = 3 *S. purpuratus* (test diameter ≥ 3 in) as detailed in section 2.3. The volume was immediately split evenly into two sterilized borosilicate glass dishes (50 mm diameter, Supertek, #19.112.0050). Treated samples received 1,000 mJ/cm^2^ UVB radiation as described above. Samples were collected into 15 mL sterile Falcon tubes and recovered for 6 h *in vitro* at 17°C, in the dark. Cells were pelleted (7,197 x g, 5 min, 17°C) and the supernatant was discarded. Cells were washed by gently resuspending the cell pellet in 1 mL of cold anticoagulant (CMFSW-E) by pipetting before pelleting (10,000 x g, 5 min, 17°C) and discarding the supernatant. Cell pellets were stored at -80°C until proceeding with lysis. Cell pellets were lysed by resuspending in 1x cell lysis buffer (Cell Signaling Technology #9803) and sonicating on ice (5 sec on, 30 sec off, 50% amplitude) using a Qsonica ultrasonic processor. Samples were centrifuged at 16,000 x g for 10 min at 4°C. The supernatant (crude protein extract) was recovered, aliquoted into sterile tubes on ice, and stored at -80°C. Protein concentration was determined with a Pierce™ Bradford Plus Protein Assay Kit and BSA standard curve (Thermo Fisher Scientific, #23236). Sample mastermixes consisting of 15 µg protein extract, 1x dithiothreitol (DTT) (Cell Signaling Technology, #14265), and 1x Blue Loading Buffer (Cell Signaling Technology, #56036) were incubated at 95°C for 5 min before loading into a 4-20% Mini-PROTEAN TGX precast SDS-PAGE gel (BioRad, #4561096). Blue prestained protein marker (11 – 250 kDa, Cell Signaling Technology, #59329) was also loaded (3.5 μL). Electrophoresis was performed at 90 V for 10 min, then at 120 V for 1 h using a Mini-PROTEAN Tetra vertical electrophoresis cell (BioRad) with 1x Tris-Glycine SDS-PAGE running buffer (25 mM Tris, 192 mM glycine, 0.1% SDS, pH 8.3). Proteins were transferred to 0.2 μm pore size nitrocellulose membranes (Cell Signaling Technology, #12369) with 1x Tris-Glycine transfer buffer (25 mM Tris, 192 mM glycine, 20% (v/v) methanol, pH 8.3) in a BioRad mini-PROTEAN tetra electrophoresis cell at 100 V for 1 h at 4°C. Membranes were transiently stained to confirm protein transfer with MemCode™ reversible protein stain kit (Thermo Scientific, #24580). Membranes were washed with Milli-Q water and blocked with 5% (w/v) non-fat dry milk in 1x Tris-buffered saline with 0.1% Tween-20 (TBST) for 1 h at room temperature with slow rocking. Membranes were washed three times with 1x TBST and incubated overnight in primary antibody solutions (1:1000 dilutions in 1x TBST with 5% w/v BSA) at 4°C with slow rocking. Primary antibodies included ubiquitin rabbit polyclonal antibody (Cell Signaling Technology, #58395), phospho-ATM/ATR Substrate Motif (Cell Signaling Technology, #6966), phospho-Akt Substrate Motif (Cell Signaling Technology, mix of #9614 and #10001), phospho-MAPK/CDK Substrate Motif (Cell Signaling Technology, mix of #9477 and #2325), and phospho-PKC Substrate Motif (Cell Signaling Technology, #6967). β-actin rabbit polyclonal antibody was used as a loading control (Cell Signaling Technology, #4967, diluted 1:1000 in 1x TBST with 5% w/v BSA). Membranes were washed three times with 1x TBST before incubating with HRP-conjugated secondary antibodies (Cell Signaling Technology anti-remazol blue mouse mAb HRP conjugate #46387 [dilution, 1:2000], used to detect the protein ladder, and anti-rabbit IgG HRP-liked antibody #7074 [dilution, 1:3000]) for 1 h at room temperature. Membranes were washed three times with 1x TBST. HRP signal was visualized using SignalFire (Cell Signaling Technology, #6883). Membranes were imaged using a UVP ChemStudio imaging system (Analytik Jena). Western blot images (8 bit grayscale, PNG format) were quantified using FIJI (ImageJ, NIH, v.1.54p). Regions of interest (ROIs) were drawn to encompass signal in bands (for β-actin loading controls) or over entire lanes (for PTM-motif substrates). A ROI of similar dimensions was also drawn to capture a section of the blot in which no protein was loaded for background correction. The “measure” function within the ROI manager was used to extract the Raw Integrated Density (RawIntDen) values for each band representing total pixel intensity within the ROI. ROI intensities for each band or lane were first background corrected by subtracting the RawIntDen value of the corresponding background ROI. Next, the corrected RawIntDen values were normalized to the background-corrected RawIntDen value of the β-actin loading control band for each sample. Normalized integrated density values from 6 biological replicates per treatment (n = 6 control, n = 6 UVB-treated) were analyzed in Microsoft Excel using one-tailed paired t-tests to assess statistical significance. Significance was set to *p* < 0.05.

## 3 Results

### 3.1 Sea urchin coelomocytes are highly resistant to UVB-induced DNA damage

We selected the purple sea urchin (*S. purpuratus*) for these studies due to its longevity (lifespan > 50 years), absence of reported neoplasia, high-quality reference genome, and well-characterized immune system (15,28,72,73). To evaluate the DNA damage response and linkage to immune activation in sea urchin coelomocytes, we initiated the response using UVB radiation as the genotoxic agent due to its well-known mechanism of action. UVB creates bulky, helix-distorting DNA lesions, including cyclobutane pyrimidine dimers and pyrimidine-pyrimidone photoproducts, which are repaired via the nucleotide excision repair (NER) pathway (74,75). In addition, UVB allowed for precise dosing without washing steps required to remove chemical mutagens. UVB is also a biologically relevant stressor of sea urchins, which begin their lifecycle as planktonic embryos and larvae in the sunlit photic zone of marine systems. Consequently, sea urchin genomes encode a repertoire of DNA repair mechanisms necessary to combat UV-induced damage (76,77).

Coelomocytes were collected from three adult *S. purpuratus*, exposed to increasing doses of UVB radiation, and allowed to recover for 24 h *in vitro.* Coelomocytes tolerated high doses of UVB, with an LD_50_ value of 7232 mJ/cm^2^ UVB and 91% viability following exposure to 1000 mJ/cm^2^ (Fig. 1a). In contrast, human peripheral blood mononuclear cells (PBMCs; ATCC PCS-800-011) demonstrated a sharp decline in viability following exposure to UVB with an LD_50_ value of 871 mJ/cm^2^. Because UVB exposure induces DNA strand breaks indirectly during the repair process (74,75), we therefore used the alkaline comet assay (single-cell gel electrophoresis) to quantify DNA damage resulting from UVB exposure, which is a sensitive technique to measure DNA strand breaks at the single cell level. Damaged DNA migrates out of the nucleus under electrophoresis, forming a “comet tail” that is visualized after applying a fluorescent DNA stain. (78,79) (Fig. 1b). DNA damage is quantified as the percentage of DNA in the comet tail (“tail DNA percent”), which is proportional to the extent of DNA fragmentation. At all assayed timepoints (1, 6, and 24 hours post-UVB exposure), coelomocytes treated with 1000 mJ/cm^2^ UVB exhibited significantly higher average tail DNA percentages and a larger fraction of damaged cells compared to controls (one-way ANOVA with post-hoc Tukey’s test; *p* < 0.01) (Fig. 1c,d). Based on these results, a dose of 1000 mJ/cm^2^ was chosen for subsequent experiments because it resulted in high levels of DNA damage while causing low levels of cell death over 24 h.

**Figure 1.**
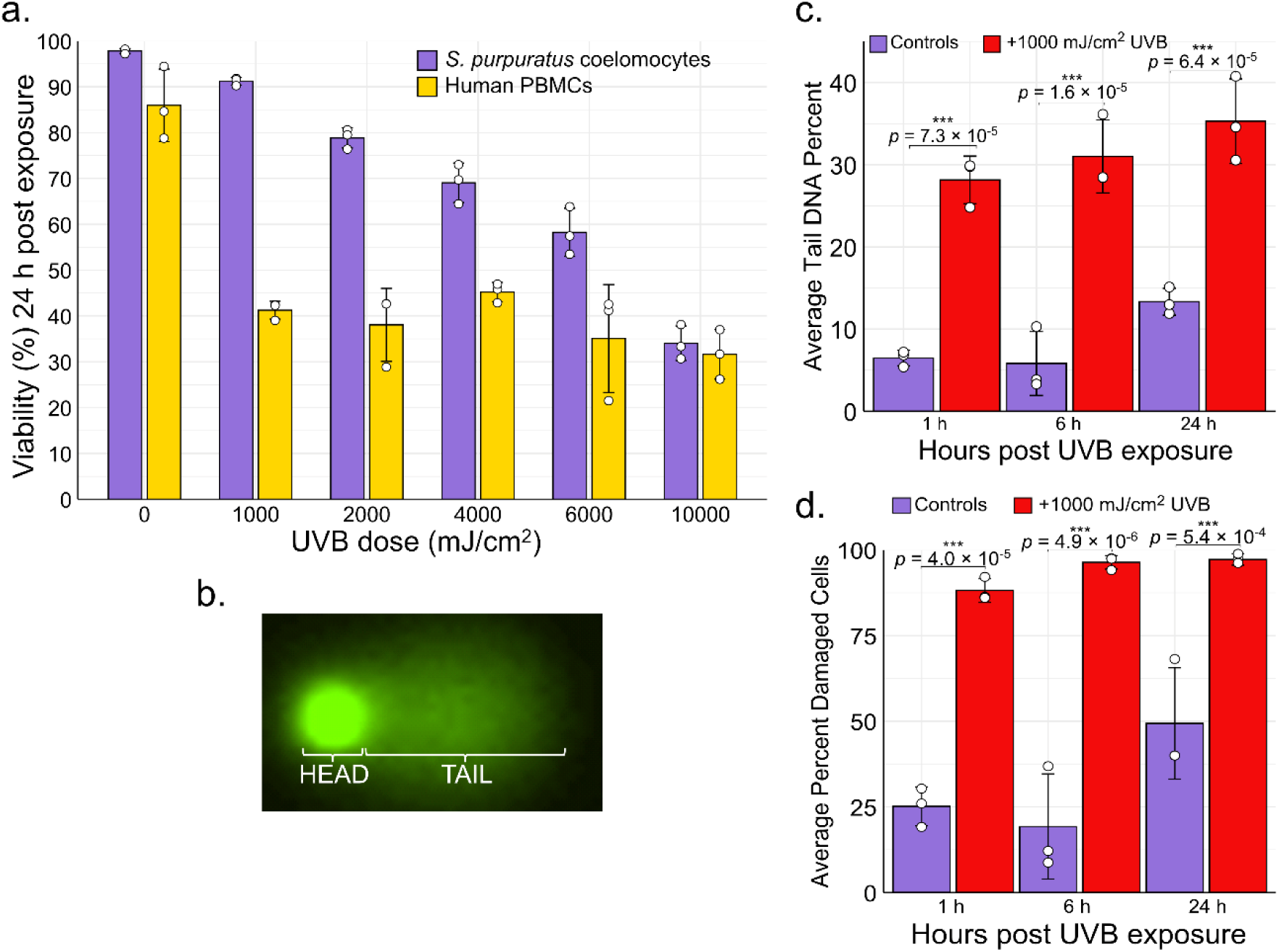
UVB-induced DNA damage and viability in sea urchin coelomocytes and human PBMCs. (a) Viability of *S. purpuratus* coelomocytes and human peripheral blood mononuclear cells (PBMCs) measured 24 h after exposure to a range of UVB doses. For coelomocytes, mean values ± the standard deviation of biological triplicates (n = 3) are shown. For PBMCs, mean values ± the standard deviation of technical triplicates are shown. Individual points are overlaid as white circles. (b) Example of a comet with the head and tail region indicated. (c) Average tail DNA percent and (d) average percent of damaged cells (the percent of cells with any amount of DNA damage) in *S. purpuratus* coelomocytes challenged with 1,000 mJ/cm^2^ UVB at 1, 6, and 24 h post-exposure (red bars) versus control, untreated coelomocytes (purple bars). Error bars are mean values ± the standard deviation of biological triplicates (n = 3). Individual points are overlaid as white circles. Significant differences between control and UVB-treated cells were analyzed using a one-way ANOVA and post-hoc TukeyHSD test (***, adjusted *p* value < 1 x 10^-3^).

### 3.2 UVB induces rapid transcriptional changes in coelomocytes

To characterize the coelomocyte transcriptomic response to UVB challenge, bulk RNA sequencing was performed at 0, 1, 3, 6, and 24 h post-exposure (1000 mJ/cm^2^). A total of 2,581 unique genes were differentially expressed (log_2_ fold-change > 1 or < -1, Benjamini-Hochberg adjusted *p* < 0.05) and were grouped into two k-means clusters (Fig. 2a). Cluster 1 included 1,247 genes that were significantly upregulated in UVB-treated cells compared to controls, while cluster 2 included 1,334 genes that were significantly downregulated in UVB-treated cells compared to controls (Fig. 2b,c). A two-way ANOVA was used to assess the effects of treatment, timepoint, and their interaction on gene expression within each cluster. Treatment had a significant effect on gene expression in both clusters (*p* = 6.2 × 10 ¹³ for cluster 1, *p* = 1.4 × 10^-8^ for cluster 2), indicating a robust transcriptional response to the experimental condition. In contrast, the treatment × timepoint interaction was not significant for either cluster (*p* = 0.14 for cluster 1, *p* = 0.40 for cluster 2), suggesting that the effect of treatment was consistent across timepoints. Timepoint alone was not a significant factor for cluster 1 (*p* = 0.39) but was significant for cluster 2 (*p* = 1.1 × 10^-4^), indicating that genes in cluster 2 exhibited time-dependent expression changes independent of treatment. These results indicate that treatment was the primary determinant of expression differences in both clusters.

**Figure 2.**
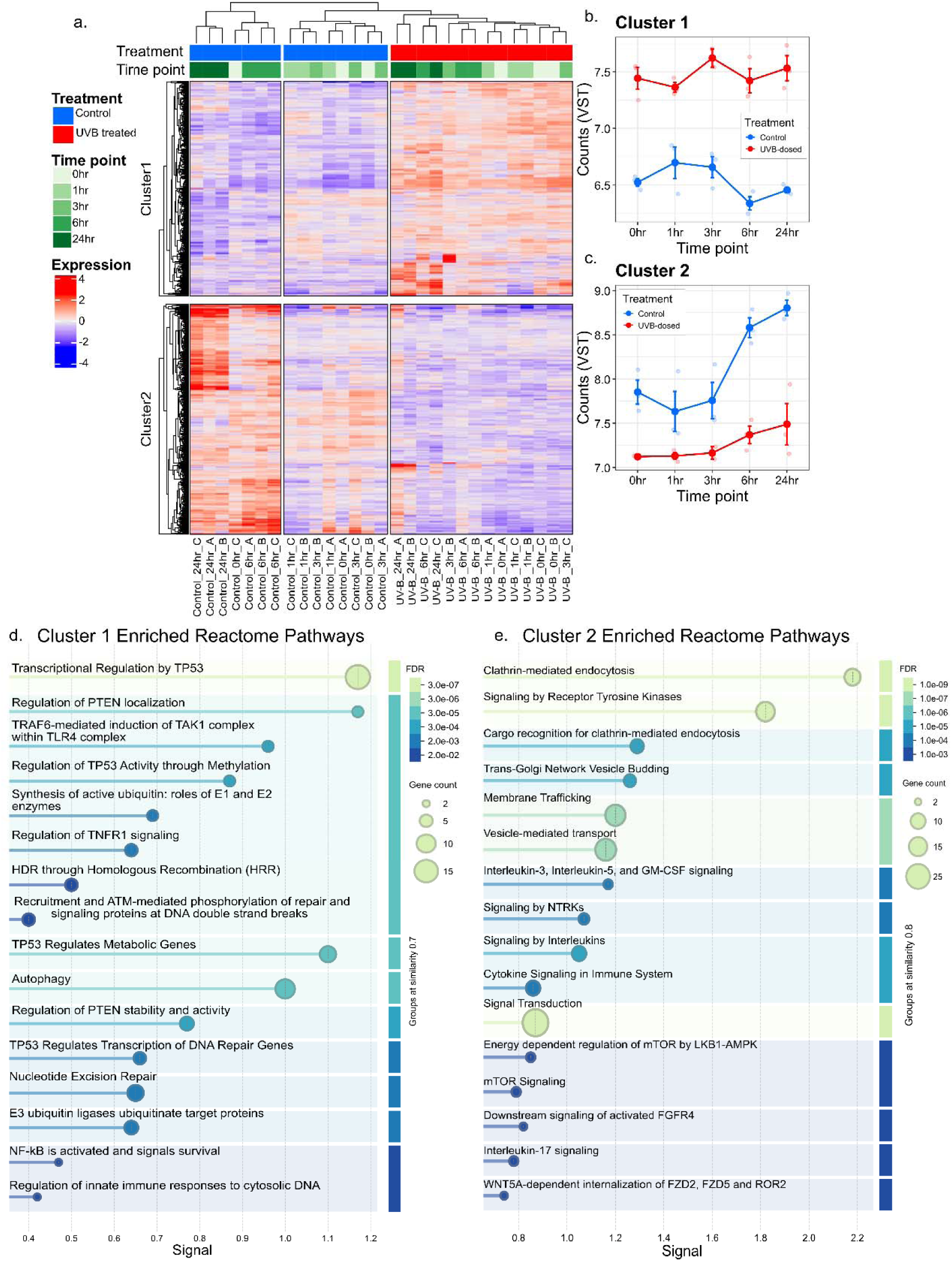
Bulk RNA sequencing revealed transcriptional changes in coelomocytes following UVB treatment. (a) The heatmap displays the expression levels of 2,581 differentially expressed genes (adjusted *p* value < 0.05, |log_2_foldchange| > 1) comparing UVB-treated samples to controls. Rows represent individual genes clustered into two groups (Cluster 1 and Cluster 2) based on k-means clustering. Columns represent samples grouped by treatment (control or UVB-treated) and time point (0, 1, 3, 6, and 24 h). The color scale indicates centered, variance-stabilized transformed (VST) gene expression values representing upregulation (red), downregulation (blue), or no change (white). Annotations indicate treatment and timepoint, with a timepoint gradient from 0 h (light green) to 24 h (dark green) for UVB-treated samples (red), and control samples (blue). X-axis labels in (a) denote the sample name in treatment_timepoint_replicate format. Mean VST gene expression counts for (b) genes in Cluster 1 and (c) genes in Cluster 2 are shown across all time points comparing treatments. Each small circle represents the mean expression of all genes in a cluster from an individual biological replicate (n = 3 replicates per treatment per time point). Group means are shown as larger points that are interconnected with a line to visualize temporal trends. Error bars represent ±1 standard error of the mean (±1 SEM) calculated from biological replicates (n = 3) per treatment per time point. (d) Selected enriched Reactome pathways ascribed to the largest protein-protein-interaction (PPI) network encoded by Cluster 1 (upregulated) genes. (e) Selected enriched Reactome pathways ascribed to the largest PPI network encoded by Cluster 2 (downregulated) genes. The x-axis in (d) and (e), “signal”, is a STRING enrichment strength score for each Reactome pathway. Enriched terms in (d) and (e) are colored according to the false discovery rate (FDR). Complete lists of enriched terms are reported in Supplementary Table 1.

To understand which cellular pathways were up- and down-regulated in coelomocytes in response to UVB challenge, we performed pathway enrichment analysis using STRING with highest confidence settings (68). Analysis of protein sequences corresponding to upregulated genes (Cluster 1) revealed a large protein-protein-interaction network characterized by 74 GO Biological Process terms, 17 GO Molecular Function terms, 68 GO Cellular Component terms, 8 KEGG pathways, and 190 Reactome pathways (80) (Supplementary Table 1). The GO and KEGG terms were related to ribosome biogenesis, mitochondrial oxidative phosphorylation, and nucleotide excision repair. Enriched Reactome pathways included DNA repair, autophagy, ubiquitination, pathways involving tumor suppressors p53 and PTEN, the early DNA-damage response involving ATM, and pathways involving immune signaling molecules such as NF-κB, TRAF, and TNFR (Fig. 2d). Notably, enriched terms describing apoptotic pathways were not observed in this bulk RNAseq analysis.

There was a shared signature of 90 upregulated genes across all time points which included ribosomal, mitochondrial oxidative phosphorylation, and ubiquitin-related genes (Supplementary Table 1). The increased expression of oxidative phosphorylation and ribosomal pathways over 24 h post-UVB treatment is consistent with previous reports using other cell types and model systems (81,82), illustrating a fundamental link between cellular stress and energy metabolism driven by the energetic cost of DNA damage repair (82). Two genes upregulated across all time points encode the ubiquitin-ribosomal fusion proteins UBA80 (RPS27A) and UBA52 (RPL40). Notably, 14 of the significantly enriched Reactome pathways describing Cluster 1 genes, including “NF-κB is activated and signals survival” and “Regulation of innate immune responses to cytosolic DNA”, were flagged as significant solely due to these two genes (Supplementary Table 1).

Pathway enrichment analysis of protein sequences corresponding to downregulated genes (Cluster 2) revealed a large protein-protein-interaction network characterized by 40 GO Biological Process terms, 24 GO Molecular Function terms, 23 GO Cellular Component terms, 5 KEGG pathways, and 64 Reactome pathways (Supplementary Table 1). Notably, many of these pathways were related to endocytosis, vesicle-mediated transport, and signal transduction pathways including Wnt, mTOR, and cytokine signaling (Fig. 2e, Supplementary Table 1). Only seven downregulated genes were shared across all time points, with functions in protein trafficking, biosynthesis, G-protein signaling, and cytoskeletal reorganization (Supplementary Table 1). It is noteworthy that the 0 h timepoint showed a significant difference between UVB-treated samples and untreated controls (Fig. 2b,c). This may be related to processing that took about 15 minutes between sample collection and cell pellet freezing, allowing time for an initial transcriptomic response to UVB treatment. This rapid response may involve mRNA stabilization, as it has been previously reported that mRNA stability was the predominant means of increasing the abundance of about one quarter of UV-induced mRNAs (83,84). In particular, UVB-induced genotoxic stress has been shown to stabilize ubiquitin-encoding mRNAs (85). Notably, the number of differentially expressed transcripts initially decreased from 665 at 0 h to 288 at 1 h and 339 at 3 h post-UVB exposure, before increasing to 1101 and 1420 at 6 h and 24 h, respectively (Supplementary Table 1).

### 3.3 Single-cell RNA sequencing revealed distinct responses to UVB treatment by different coelomocyte cell types

The response of specific coelomocyte cell types to DNA damage has not been investigated previously. To address this knowledge gap, we investigated the transcriptional response of different coelomocyte cell types to genotoxic challenge using single-cell RNA sequencing (scRNA-seq) at 6 h post-UVB exposure (1000 mJ/cm^2^). This timepoint was chosen based on the strong upregulated transcriptional signature observed in the bulk RNA-seq data at 6 h of recovery (Supplementary Table 1; Fig. 2). Furthermore, prior studies have shown that both signal transduction pathways and transcription factor nuclear translocation are activated within 6 h of UV irradiation in mammalian cells (84,86). UVB-treated coelomocytes and controls (three biological replicates per condition, totaling 74,980 cells after preprocessing; Supplementary Table 2) were visualized using leiden clustering (87), revealing 22 cell clusters (Fig. 3a). Distinct separation of UVB-treated coelomocytes and controls in UMAP space was observed (Fig. 3b), demonstrating that UVB-treated cells were transcriptionally distinct from control cells. Cell clusters 0, 1, 8, 11, 17, 18, 20, and 21 were predominantly comprised of control cells (> 80%), while cell clusters 2, 4, 5, 6, 9, 12, and 15 were predominantly comprised of UVB-treated cells (> 80%). Cell clusters 3, 7, 10, 13, 14, 16 and 19 were comprised of a mix of UVB-treated and control cells (Supplementary Table 3). To determine the transcriptional response of each cell type to UVB challenge, each Leiden cluster was first identified as a coelomocyte cell type (phagocytes, red spherule cells, colorless spherule cells, or vibratile cells; Fig. 3c) based on significant upregulated expression (Benjamini-Hochberg corrected *p* < 0.05, log_2_ fold-change > 0.5) of genes ascribed to each cell type in the literature (Supplementary Table 4). We identified 15 clusters as phagocytes based on significant expression of genes belonging to the *SpTransformer* family (*SpTrf*; formerly *Sp185/333*) (88), the Toll-like receptor (*TLR*) family (30), complement component gene *SpC3* (Smith et al. 2001, 2023) and *Sp-B7L3* (89) (Supplementary Table 4). Two cell clusters (13 and 16) were identified as red spherule cells based on significant expression of the marker gene polyketide synthase 1 (*pks1*). Pks1 is key to synthesizing echinochrome A, the pigment that gives these cells their red color (90–92). Definitive marker genes for vibratile cells have not been described. However, these cells possess a single, whip-like flagellum powered by the motor protein dynein, particularly axonemal dynein (72). Therefore, significant expression of dynein heavy chain genes was used to characterize cell clusters 7, 19, and 21 as vibratile cells (Supplementary Table 4). Amassin is a protein in the coelomic fluid that mediates cell-cell adhesion by forming disulfide-bonded aggregates, forming a protein clot that captures coelomocytes (93). Interestingly, we noted significant expression (> 2 log_2_ fold-change) of amassin genes exclusively in vibratile cell clusters 7 and 19 (Supplementary Table 4), suggesting that vibratile cells play a role in the clotting reaction. Upregulated expression of *Sp-FoxJ1* and *Sp-P2rx4* was previously observed within a mix of vibratile cells and colorless spherule cells (these cell types appeared in the same density gradient fraction, thus they were not separated) (89). Consistently, significant expression of *Sp-P2rx4* was observed in all three vibratile cell clusters (7, 19, and 21) while *Sp-FoxJ1* expression was observed in two of these clusters (7 and 19) (Supplementary Table 4). As with vibratile cells, no definitive marker genes for colorless spherule cells have been described, though these cells are known to contribute to cytotoxic activity (94). A combined signature of significant expression of bactericidal permeability-increasing protein, lysozyme, and strongylocin, and non-significant expression of *Pks1* and phagocyte makers, was therefore used to identify clusters 10 and 14 as colorless spherule cells (Fig. 3c, Supplementary Table 4). Marker gene expression therefore enabled assignment of each cell cluster to a defined coelomocyte subtype.

**Figure 3.**
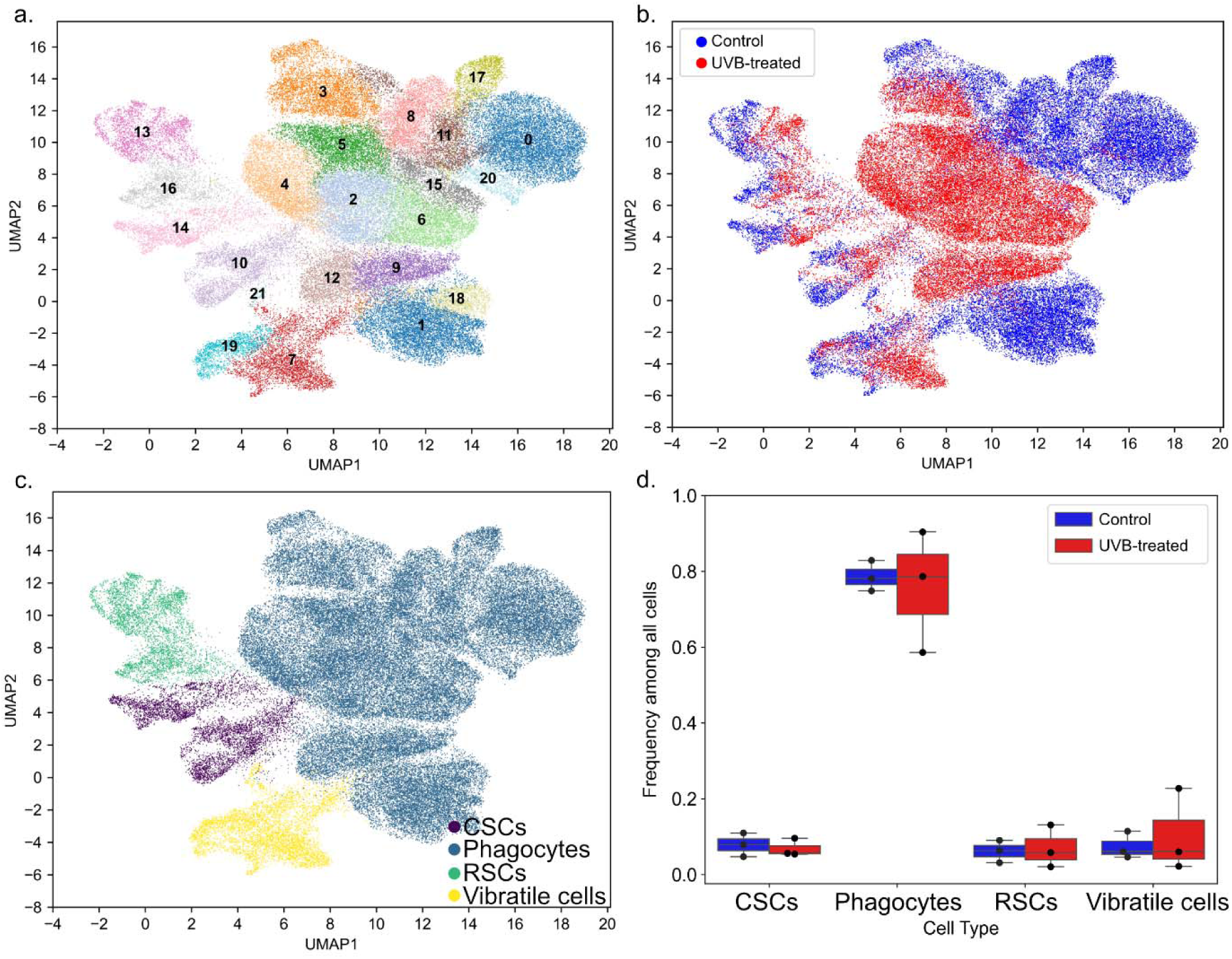
The cellular landscape of control and UVB-treated coelomocytes at 6 h recovery *in vitro* was revealed by single-cell RNA sequencing. (a) Major scRNA-seq clusters (22). (b) Control (blue) vs UVB-treated samples (red) with the same embedding as in (a). (c) Respective cell-type assignments based on marker gene expression with the same embedding as in (a). Clusters were identified as colorless spherule cells (CSCs, purple), phagocytes (blue), red spherule cells (RSCs, green), and vibratile cells (yellow). (d) Fraction of major cell types in control (n = 3, blue) and UVB-treated (n = 3, red) coelomocyte samples. The box plots indicate the median, 25^th^ and 75^th^ percentiles are the lower and upper edges of the boxes, and the whiskers show the most extreme points that do not exceed ±1.5x the interquartile range (IQR).

Cell type fractions within each sample were next explored. UVB-treated samples were comprised of 76% ± 13% phagocytes, 10% ± 9% vibratile cells, 7% ± 5% red spherule cells, and 7% ± 2% colorless spherule cells, while control samples were comprised of 79% ± 3% phagocytes, 7% ± 3% vibratile cells, 6% ± 2% red spherule cells, and 8% ± 3% colorless spherule cells (averages ± standard deviation of biological triplicates; Fig. 3d, Supplementary Table 3). No significant differences in any cell type fraction were observed when comparing UVB-treated samples and control samples (two-tailed paired *t*-test, *p* > 0.05 for all comparisons). These cell type fractions were consistent with our direct observations during cell counting and with prior observations in *S. purpuratus,* a species in which phagocytes typically represent the most abundant cell type (∼40 – 80%), followed by vibratile cells (11.9 – 20%), red spherule cells (7 – 40%), and colorless spherule cells (3.7 – 25%) (14).

Differential gene expression analysis was first conducted on the scRNA-seq data to examine transcriptomic differences in UVB-treated coelomocytes and controls, inclusive of all cell types. A total of 561 genes were significantly upregulated (*p* < 0.05 and log_2_ fold-change > 1) and 319 genes were significantly downregulated (*p* < 0.05 and log_2_ fold-change < -1) in UVB-treated cells compared to controls (Fig. 4a, Supplementary Table 5). Consistent with the bulk RNAseq data, genes involved in DNA repair and ubiquitination (particularly E3 ubiquitin ligase tripartite motif (TRIM)-like genes) were among the most highly upregulated genes in UVB-treated cells, while *wnt-1* was among those most downregulated (Fig. 4b). GO pathway analysis of these upregulated genes revealed six significant (*p* < 0.05) terms including the GO Molecular Function terms ‘zinc ion binding’ (123 genes), ‘ubiquitin-protein transferase activity’ (94 genes), and ‘DNA-binding transcription factor activity’ (17 genes); the GO Cellular Component terms ‘nucleoplasm’ (86 genes) and ‘chromatin’ (84 genes); and the GO Biological Process term ‘positive regulation of DNA-templated transcription’ (82 genes) (Fig. 4c; Supplementary Table 5). GO pathway analysis of the downregulated genes revealed twelve significant (*p* < 0.05) terms including the GO Molecular Function terms ‘metalloendopeptidase activity’, ‘growth factor activity’, ‘cytokine activity’, ‘metalloendopeptidase inhibitor activity’, ‘extracellular matrix binding’, and ‘collagen binding’; the GO Cellular Component terms ‘extracellular space’, ‘extracellular region’, and ‘extracellular matrix’; and the GO Biological Process terms ‘extracellular matrix organization’, ‘collagen catabolic process’, and ‘Smad protein signal transduction’ (Fig. 4d; Supplementary Table 5). These findings indicate that UVB challenge caused coelomocytes to suppress growth and trafficking-related functions while prioritizing DNA repair and stress response, consistent with our bulk RNA-seq observations.

**Figure 4.**
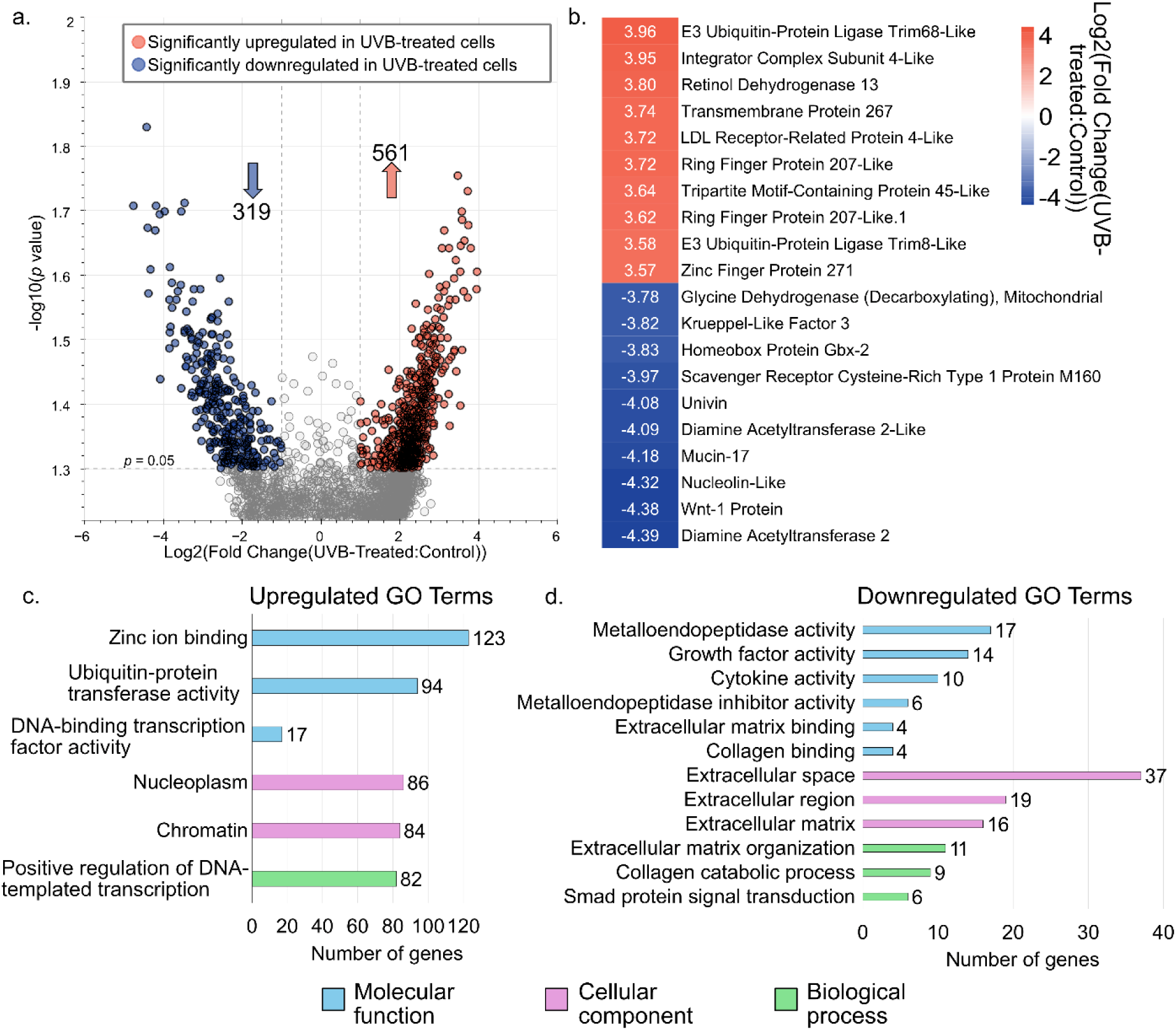
UVB-treated coelomocytes upregulate DNA repair and stress response genes while downregulating growth factors. (a) The volcano plot shows the distribution of differentially expressed *S. purpuratus* transcripts plotted according to *p* value and log_2_fold-change comparing all cell types in UVB-treated samples vs. control samples. Dashed lines indicate significance level cutoffs at *p* < 0.05 and |log_2_(fold change)| > 1. Arrows indicate the number of upregulated (orange) and downregulated (blue) genes. (b) Heatmap showing the top ten most significantly upregulated (orange) and downregulated (blue) genes comparing UVB-treated samples and controls by log_2_foldchange. (c) Bar chart of enriched GO terms representing the significantly upregulated genes in UVB-treated cells. (d) Bar chart of enriched GO terms representing the significantly downregulated genes in UVB-treated cells. All significant GO terms are shown (*p* < 0.05; Supplementary Table 5). The number of genes mapped to a particular GO term is shown on the x axis. GO = Gene Ontology.

We next used the scRNA-seq data to perform differential gene expression analysis comparing the effect of UVB treatment at the cell-type level. This cell-type-resolved approach revealed that phagocytes and vibratile cells mounted a more robust transcriptional response to UV exposure compared to red spherule cells and colorless spherule cells, with 1079, 820, 165, and 125 significantly upregulated genes (*p* < 0.05, log_2_ fold-change > 1) and 371, 147, 117, and 179 significantly downregulated genes (*p* < 0.05, log_2_ fold-change < -1) in UVB-treated phagocytes, vibratile cells, red spherule cells, and colorless spherule cells, respectively (Fig. 5a-d, Supplementary Table 6). To assess the function of the significantly upregulated and downregulated genes, the protein sequences encoded by these genes were used as input for STRING analysis under high confidence settings. For UVB-treated phagocytes, the largest protein-protein-interaction (PPI) network among upregulated gene products following k-means clustering consisted of 40 nodes (PPI enrichment *p* < 1.41 x 10^-14^) (Fig. 6a). Significantly enriched Reactome pathways describing this PPI network included terms involving tumor suppressor p53 and DNA damage sensing and repair pathways such as ‘Nonhomologous End-Joining (NHEJ)’, ‘Nucleotide Excision Repair,’ ‘Recruitment and ATM-mediated phosphorylation of repair and signaling proteins at DNA double strand breaks’, and ‘Homology Directed Repair’ (Fig. 6b; Supplementary Table 7). The 371 downregulated gene products formed nine minimal PPI networks following k-means clustering (all with 10 nodes or less). Significantly enriched pathways describing the three largest networks indicated downregulation of heat shock proteins, transcription, protein processing, and ciliary movement (Supplementary Table 7). Pathway enrichment analysis of the protein sequences encoded by the 820 significantly upregulated genes in UVB-treated vibratile cells revealed a PPI network of 37 nodes (PPI enrichment *p* value < 1.0 x 10^-16^) (Fig. 6c). Significantly enriched Reactome pathways described cell cycle regulation, DNA repair, and p53 pathways (Fig. 6d, Supplementary Table 7). The 147 downregulated gene products formed two minimal PPI networks (≤ three nodes) following k-means clustering. Only two significant functional enrichments were observed: the KEGG pathway “Glyoxylate and dicarboxylate metabolism” and the Local STRING Network Cluster “Mixed, incl. RUNX1 regulates transcription of genes involved in BCR signaling, and Runt domain” (Supplementary Table 7). Overall, our findings indicate that UVB exposure elicited a robust transcriptional response in phagocytes and vibratile cells involving the DNA damage response, p53 signaling, and repair-associated pathways.

**Figure 5.**
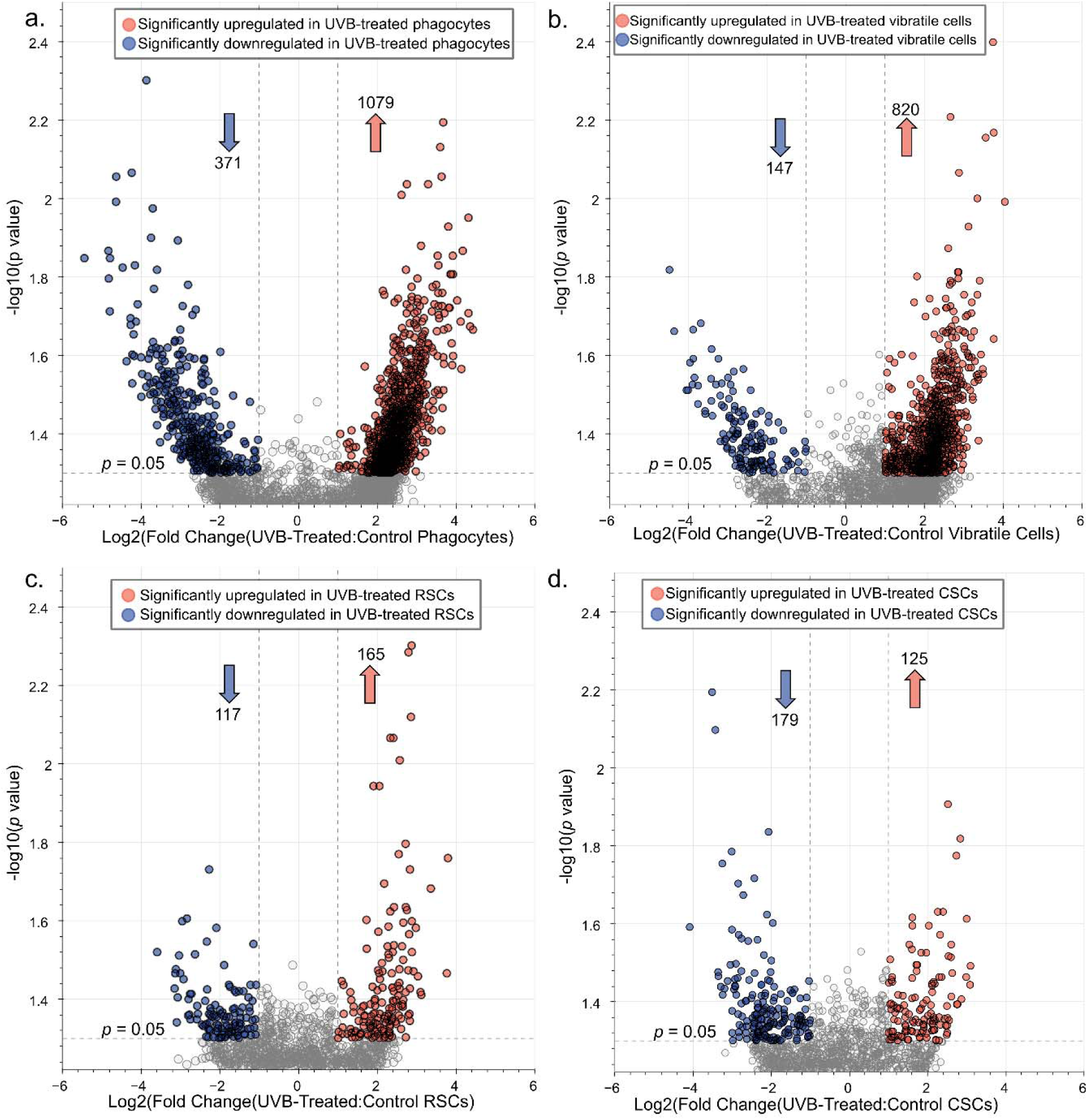
Phagocytes and vibratile cells mount a robust transcriptional response to UVB challenge, in contrast with red and colorless spherule cells. The volcano plots show the distribution of quantified *S. purpuratus* transcripts according to *p* value and log_2_fold-change comparing UVB-treated and control (a) phagocytes, (b) vibratile cells, (c) red spherule cells (RSCs), and (d) colorless spherule cells (CSCs). Dashed lines indicate significance level cutoffs at *p* < 0.05 and |log_2_(fold change)| > 1. Arrows indicate the number of upregulated (orange) and downregulated (blue) genes. Complete lists of differentially expressed genes are presented in Supplementary Table 6.

**Figure 6.**
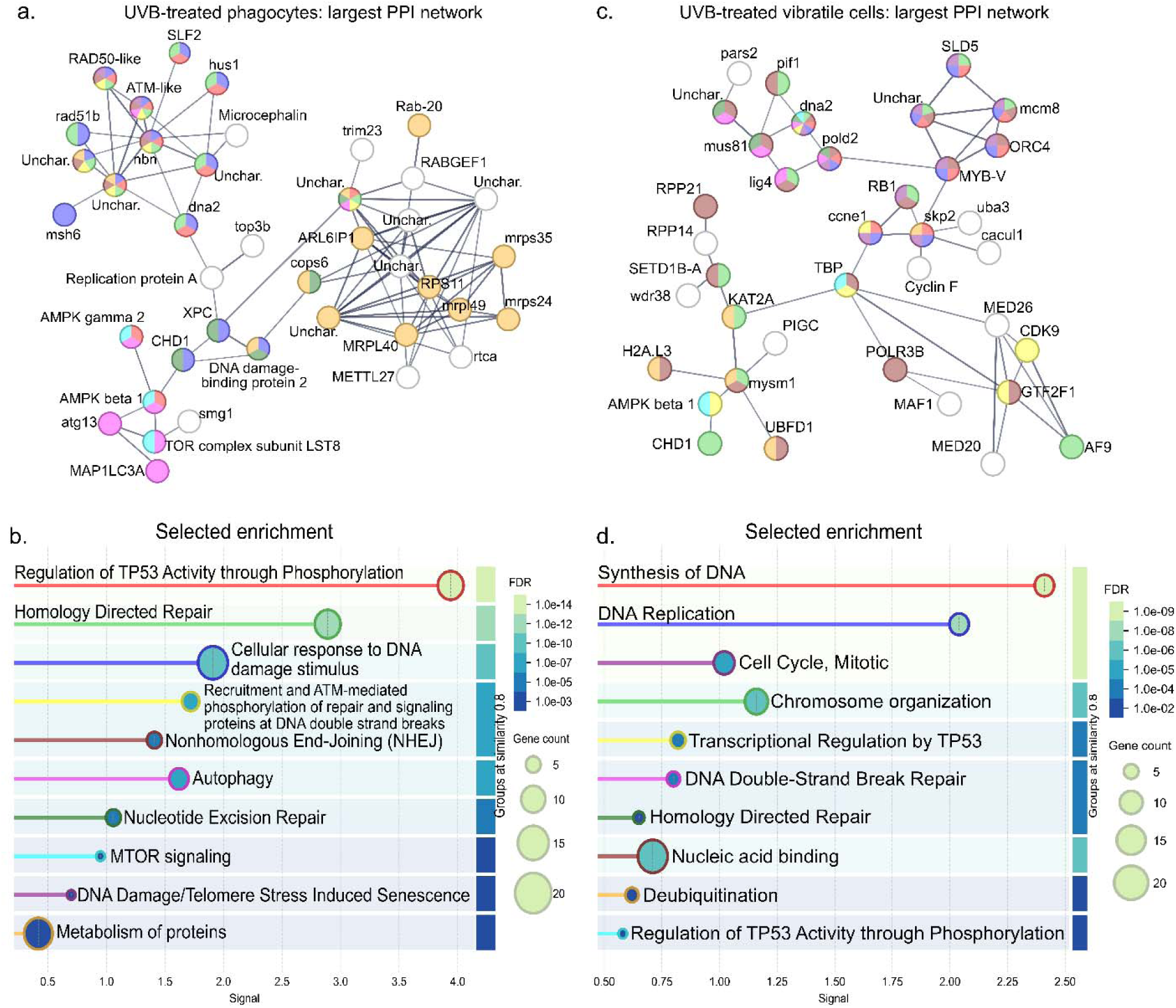
Proteins encoded by significantly upregulated genes in UVB-treated phagocytes and vibratile cells are enriched for DNA damage sensing and repair pathways. The graphical representations show the largest networks of protein-protein interactions (determined by k-means clustering) among significantly upregulated genes (*p* < 0.05, log_2_fold-change > 1) using the STRING database under high confidence settings (0.700 minimum required interaction score). (a) The largest PPI network among upregulated genes in UVB-treated phagocytes. Nodes are colored according to enriched terms shown in (b). (b) Selected enriched pathways ascribed to the phagocyte PPI network in (a). (c) The largest PPI network among significantly upregulated genes in UVB-treated vibratile cells. Nodes are colored according to enriched terms shown in (d). (d) Selected enriched pathways ascribed to the vibratile cell PPI network in (c). Enriched terms in (b) and (d) are colored according to the false discovery rate (FDR). Complete lists of enriched terms are given in Supplementary Table 7.

In contrast with phagocytes and vibratile cells, pathway enrichment analysis of protein sequences encoded by the 165 significantly upregulated genes in UVB-treated red spherule cells resulted in only minimal PPI networks. The largest PPI network (PPI enrichment *p* = 1.07 x 10^-4^) contained only seven nodes, representing the Reactome pathway ‘Endosomal Sorting Complex Required for Transport (ESCRT)’ (Supplementary Figure 1a). Other minimal networks were enriched for pathways described as ‘ribosomal protein’, ‘extrinsic apoptotic signaling pathway in absence of ligand’, and ‘inhibition of the proteolytic activity of APC/C required for the onset of anaphase by mitotic spindle checkpoint components’ (Supplementary Figure 1a, Supplementary Table 7). Interestingly, the enriched apoptotic signaling pathway consisted of only two genes, Bcl-2 homologous antagonist/killer-like (LOC100890351) and Bcl-2-like protein 1 (LOC100890351). Bcl-2 family proteins are known to induce caspase activation and regulate apoptosis (95). In red spherule cells, the 117 downregulated gene products formed only one PPI network with two nodes, describing downregulation of spliceosome activity (Supplementary Table 7). Similar to red spherule cells, analysis of protein sequences encoded by the 125 significantly upregulated genes in UVB-treated colorless cells produced only three minimal PPI networks, each with two nodes (Supplementary Figure 1b). Enriched terms describing these networks were ‘Multivesicular body sorting pathway’, ‘Cytosolic small ribosomal subunit’, and ‘DNA Damage Recognition in GG-NER’ (Supplementary Figure 1b, Supplementary Table 7). Analysis of the 179 downregulated gene products formed a two-node PPI network, though no significantly enriched pathways were observed. Of the four coelomocyte cell types, phagocytes and vibratile cells therefore mounted a robust transcriptional response to UVB challenge, while red spherule cells and colorless spherule cells demonstrated a comparatively minor response.

### 3.4 Immune genes are upregulated in response to UVB challenge

In addition to genes involved in DNA repair, p53 pathways, cell cycle arrest, and autophagy (Fig. 6b,d), UVB challenge also resulted in significant expression of various genes encoding proteins with functions in the immune system. We grouped these into five broad categories based on the echinoid immune gene repertoire: (i) Pattern Recognition Receptor (PRR), (ii) Transcription factor, (iii) Complement, (iv) Intracellular signaling, and (v) Effector (28,33) (Fig. 7; Supplementary Table 8). Overall, UVB-treated phagocytes and vibratile cells had more significantly upregulated immune genes compared to red and colorless spherule cells (Fig. 7). Transcripts of PRRs such TLRs, SRCRs, RLRs, and NLRs were most abundant in phagocytes, as were transcription factors including *TCF20, GATA, ETS, BBX,* and *ATF-2*. Effector immune genes encoding TRIM E3 ubiquitin ligases, as well as genes involved in intracellular signaling (*TRAF*, *NF-*κ*B*, *TNFR*), were represented among all cell types, although were most abundant in phagocytes. Complement component genes annotated as *C2* and *C3* were also significantly upregulated following UVB treatment. However, the *C2* gene (LOC753884) is most likely a misannotated *Bf* (factor B) gene, consistent with the lack of a classical complement pathway in sea urchins (96). Interestingly, significant upregulation of many zinc finger genes was observed among phagocytes and vibratile cells, but none were observed in red or colorless spherule cells (Fig. 7). In all categories of cell types, many significantly upregulated genes were annotated as “uncharacterized”, with higher abundances in phagocytes (304) and vibratile cells (219) compared to red spherule cells (44) and colorless spherule cells (37) (Supplementary Figure 2a). Pathway enrichment analysis of the proteins encoded by these uncharacterized genes in phagocytes revealed a significant enrichment of encoded proteins with zinc finger (KW-0863; *p* = 1.69 x 10^-9^), NACHT, (PF05729; *p* = 2.71 x 10^-12^), and RING domains (SM00184; *p* = 1.9 x 10^-3^) (Supplementary Figure 2b). Protein sequences corresponding to uncharacterized genes in vibratile cells were enriched in NACHT domains (PF05729; *p* = 7.4 x10^-4^) (Supplementary Figure 2c). No significant enrichment was detected among protein sequences encoded by uncharacterized genes in red or colorless spherule cells (Supplementary Table 9). This data indicates immune system activation concurrent with DNA repair in response to UVB challenge.

**Figure 7.**
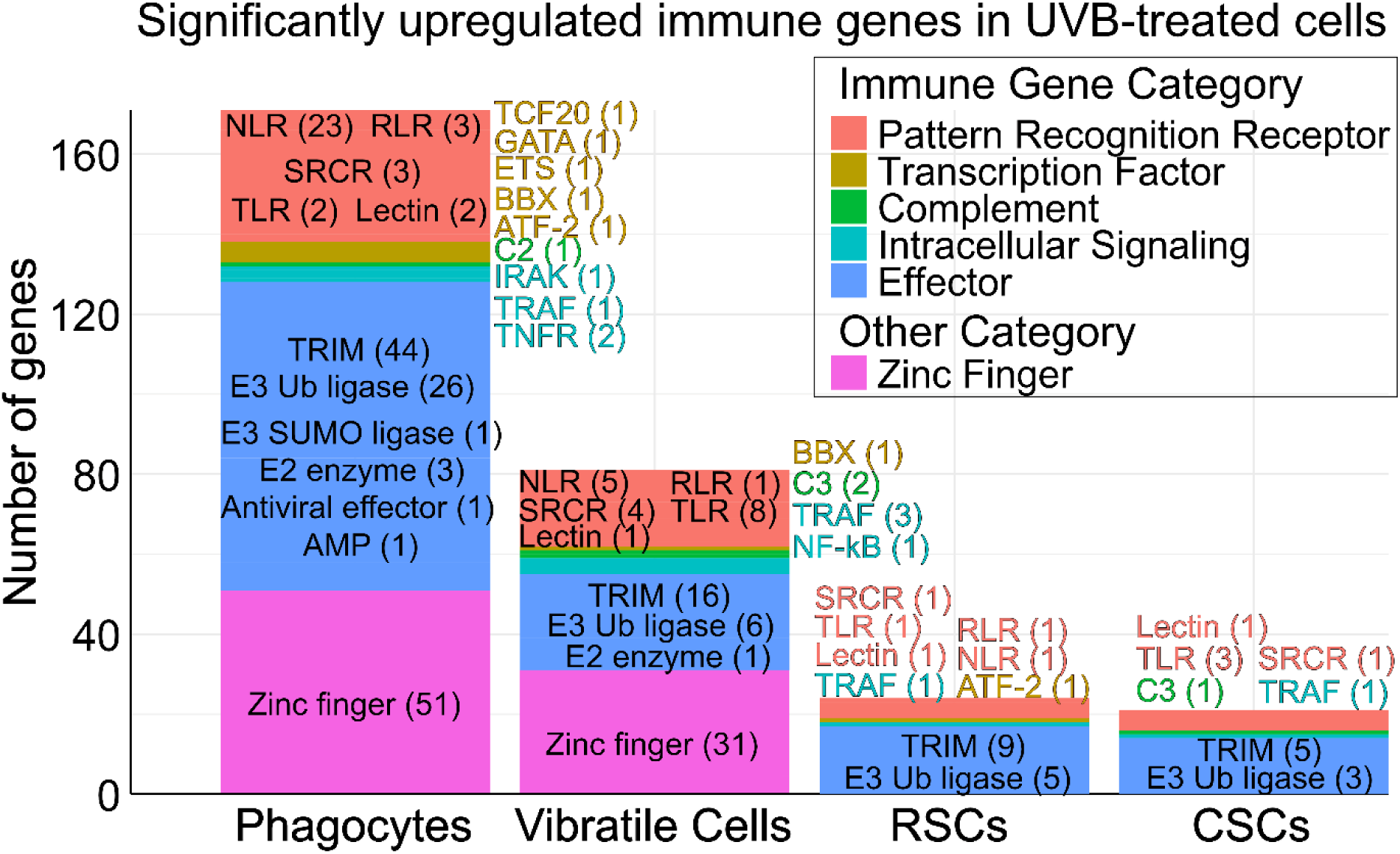
Immune system genes and zinc fingers are upregulated in response to UVB treatment in coelomocyte cell types. The stacked bar chart shows categorized groups of significantly upregulated (*p* < 0.05, log_2_ fold-change > 1) immune system genes and zinc finger genes in UVB-treated coelomocytes compared to control coelomocytes based on cell type. Examples of specific gene group names and the number of genes detected (within the parentheses) are overlaid. Complete gene lists are presented in Supplementary Table 8.

### 3.5 Functional validation of the transcriptomic data confirmed that autophagy, ubiquitination, and phosphorylation are major responses to UVB

Our transcriptomic analyses suggested that autophagy was a major response to UVB challenge in sea urchin coelomocytes (Fig. 2d), particularly in UVB-treated phagocytes (Fig. 6b). To functionally validate this observation, we quantified autophagic signal in *S. purpuratus* coelomocytes at 6 h post UVB exposure (1000 mJ/cm^2^) using the Abcam Autophagy Assay kit to quantify autophagic vacuoles. We observed a significant increase (Wilcoxon rank sum test *p* = 1.65 x 10^-4^) in autophagic fluorescent signal (FITC signal normalized to Hoechst 33342 signal) in UVB-treated cells compared to control cells (Fig. 8a,b), confirming autophagy as a major response to UVB stress.

**Figure 8.**
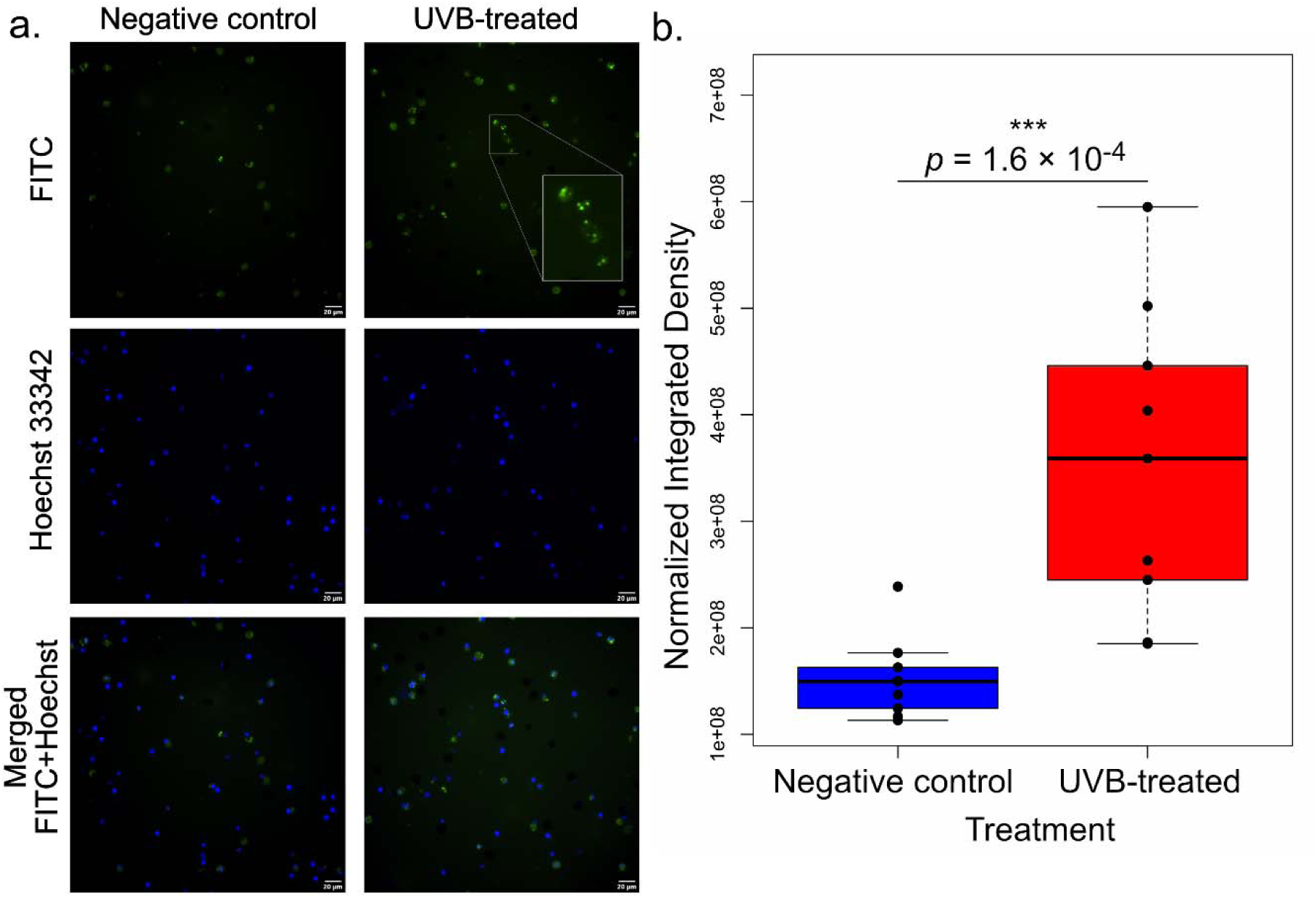
Functional validation of enhanced autophagy in UVB-treated coelomocytes. (a) Representative microscopy images of *S. purpuratus* coelomocytes subjected to no challenge (negative control) or 1000 mJ/cm^2^ UVB radiation, assayed for autophagic signal after 6 h of recovery. Autophagic vacuoles were labelled using the Abcam Autophagy Assay Kit (ab139484) (FITC signal, green), while nuclei were labelled with Hoechst 33342 stain (blue). The inset in the second microscopy image shows a digitally enlarged view of a region of interest within the FITC channel in UVB-treated cells, highlighting green puncta corresponding to labeled autophagosomes. (b) Boxplot showing the significant difference in autophagic FITC signal normalized to Hoechst signal comparing untreated and UVB-treated coelomocytes. Individual data points are overlaid as black circles (n = 3 biological replicates analyzed in technical triplicate). ***, *p* < 1 × 10^-3^ , Wilcoxon rank sum test. Scale bars are 20 μm.

Post-translational modifications (PTMs) of target proteins are known to have important functions in the DDR and innate immunity (97–101). Our transcriptomic analyses suggested that ubiquitination is a major upregulated response to UVB challenge in sea urchin coelomocytes (Fig. 2d), as indicated by upregulated expression of E3 ubiquitin ligase TRIM genes (Fig. 4b; Fig. 7). To further investigate this observation, western blot analysis with an anti-ubiquitin antibody (Cell Signaling Technology, #58395) was used to compare the ubiquitinated protein signal between crude protein extracts isolated from UVB-treated (1000 mJ/cm^2^) and control coelomocytes after 6 h of recovery *in vitro*. Results showed a significantly higher ubiquitinated protein signal (normalized to β actin loading control signal) in the UVB-treated samples compared to controls (one-tailed paired *t*-test, *p* = 0.006) (Fig. 9a,b; Supplementary Figure 3a), which was consistent with the transcriptomic findings. Our transcriptomic data also suggested increased phosphorylation was an important factor in the DNA damage response as demonstrated by the upregulation of genes involved in signaling pathways regulated by phospho-PTMs (Fig. 6b,d). We therefore conducted western blot analyses using anti-Phospho PTM substrate motif antibodies to probe conserved downstream signaling substrates associated with DNA damage and the stress response. These antibodies included phospho-ATM/ATR Substrate Motif (CST #6966), phospho-Akt Substrate Motif (CST #9614 and #10001), phospho-MAPK/CDK Substrate Motif (CST #9477 and #2325), and phospho-PKC Substrate Motif (CST #6967). While the phosphopeptide signal for ATM/ATR substrate motifs tended to be higher in UVB treated samples, there was no significant difference between UVB-treated and control samples (*p* = 0.07, one tailed paired *t*-test, n = 6) (Fig. 9c,d, Supplementary Figure 3b). In contrast, significantly increased phosphopeptide signal was observed for phospho-AKT substrate motif (*p* = 0.03), phospho-MAPK/CDK substrate motif (*p* = 0.023), and phospho-PKC substrate motif (*p* = 0.004) in UVB-treated samples compared to controls (one-tailed paired *t*-test, n = 6) (Fig. 9e-j, Supplementary Figure 3c-e). This suggested that activation of these signaling pathways had important functions in regulating cell survival and stress response in sea urchin coelomocytes following UVB challenge. This work further highlights the utility of these anti-Phospho PTM substrate motif antibodies as screening tools, even in non-traditional model organisms, and enables future follow-up PTM-proteomic studies to identify key protein regulators.

**Figure 9.**
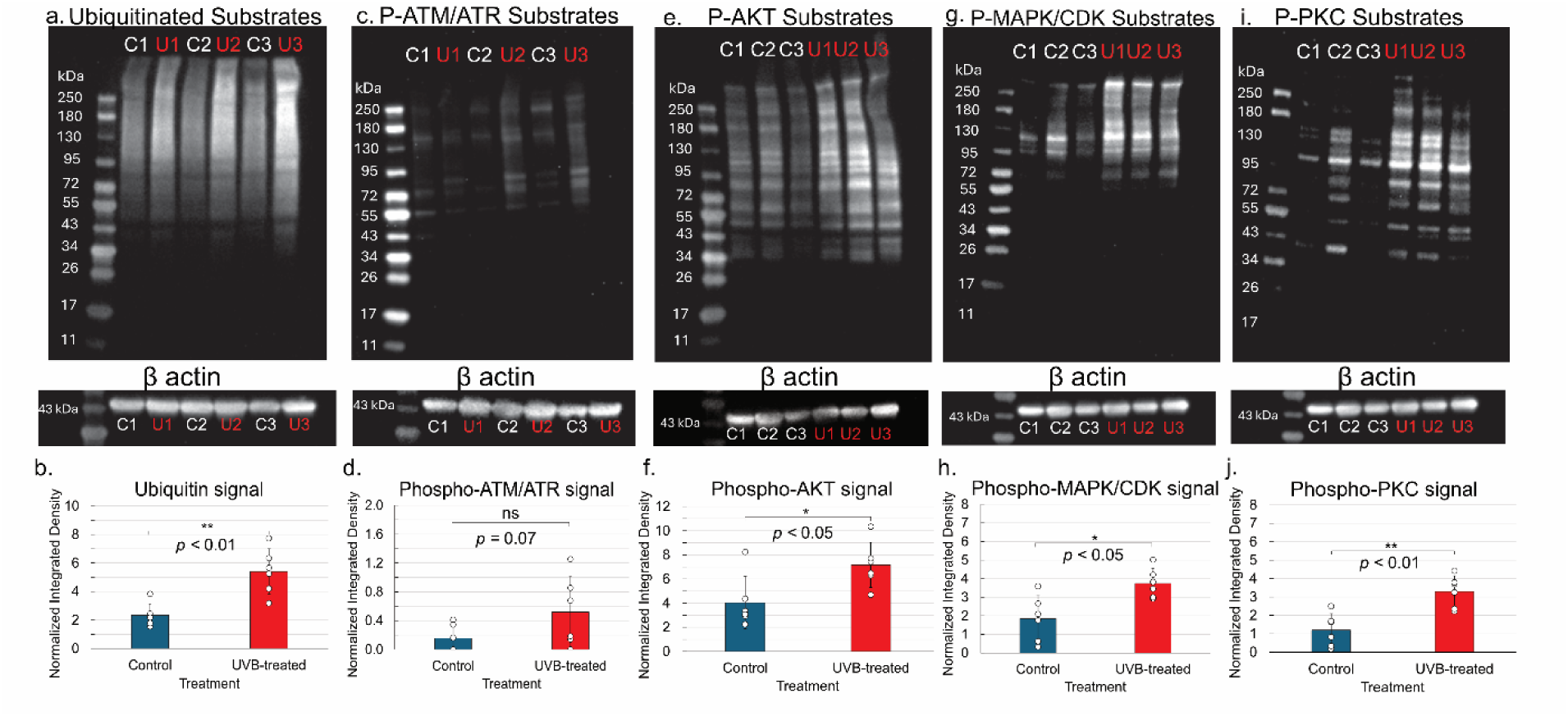
Enhanced ubiquitination and phosphorylation in UVB-treated coelomocytes was confirmed using western blot. Representative western blots showing (a) ubiquitinated, (c) phosphorylated ATM/ATR substrate, (e) phosphorylated AKT substrate, (g) phosphorylated MAPK/CDK substrate, and (i) and phosphorylated PKC substrate signal in biological triplicate controls (C, white font) and UVB-treated (U, red font) samples. Beta actin loading controls are shown below. Quantification and statistical testing (one-tailed paired t-test) of western blot signals are shown in (b), (d), (f), (h), and (j). Individual data points representing n = 6 biological replicates are overlaid as white circles. Bars represent mean values, and error bars represent ± the standard deviation of n = 6 biological replicates. * *p* ≤ 0.05, ** *p* ≤ 0.01, *** *p* ≤ 0.001. P-MAPK/CDK and P-PKC blots were run simultaneously and therefore have the same β actin loading control. Additional western blots are presented in Supplementary Figure 3.

## 4 Discussion

### 4.1 Overview of the coelomocyte transcriptional response to UVB challenge

This study revealed that immune cells of the long-lived purple sea urchin, *S. purpuratus*, mount a robust transcriptomic response to UVB-induced DNA damage, with upregulated expression of genes involved in DNA repair, autophagy, and innate immunity, and downregulation of genes involved in growth and trafficking. These findings suggest that coelomocytes suppress growth and trafficking-related functions in response to UVB challenge in favor of reallocating resources toward repair, proteostasis, and survival. Within the data generated by the timecourse of bulk-RNA sequencing, we noted two genes encoding the ubiquitin-ribosomal fusion proteins UBA80 (RPS27A) and UBA52 (RPL40) that were upregulated across all timepoints. Both of these genes have been linked to the DDR, as both are required for DNA repair by fine-tuning the spatiotemporal regulation of repair proteins at sites of DNA damage (101). The observed consistent upregulated expression of *UBA80* and *UBA52* in UVB-treated cells over 24 h implies additional functions for the encoded ribosomal proteins that respond to DNA damage, protein damage, or a combination of both. These findings are consistent with the concurrent upregulation of autophagy and ubiquitin-mediated pathways. In addition to *UBA80* and *UBA52*, upregulated genes within the bulk-RNA sequencing data were mainly comprised of other ribosomal transcripts, suggesting that the encoded ribosomal proteins may have extra-ribosomal functions. However, only a few ribosomal proteins have been demonstrated to directly participate in regulating the DDR pathway, highlighting potential unexplored ribosomal proteins with extra-ribosomal functions (102).

### 4.2 Phagocytes and vibratile cells mount a robust transcriptional response to UVB challenge, in contrast with red and colorless spherule cells

Phagocytes and vibratile cells exhibited a markedly higher number of significantly upregulated genes in response to UVB-treatment compared to red and colorless spherule cells. Interestingly, significant upregulation of many zinc finger genes was observed only among these cell types. Zinc finger proteins play an important role in immune system regulation both at the transcriptional and post-transcriptional level, and have been implicated in a broad range of processes including cytokine production, immune cell activation, immune homeostasis and antiviral innate immunity (103,104). Significant enrichment of pathways related to tumor suppressor p53 was also observed in UVB-challenged phagocytes and vibratile cells. Sea urchins possess only a single gene belonging to the p53 family of transcription factors, the ancestral family member *TP63* that is classically known for maintaining genomic integrity of the germline (105,106). Our finding that these cell subtypes activate *TP63*-related pathways in response to genotoxic stress is consistent with prior studies showing increased levels of p63 protein in sea urchin embryos exposed to ionizing radiation (107). This suggests that *TP63* has core functions in mediating DDR signaling beyond the germline in sea urchins. Functional characterization of this tumor suppressor and other tumor suppressors in sea urchins is expected to provide further insight into their role in maintaining genome stability in germline and somatic tissues.

The robust transcriptional response in phagocytes is not unexpected, as phagocytes are the most abundant coelomocyte population in sea urchins and are known to have important functions in pathogen clearance, wound healing, and immune surveillance (15,72). Their responsiveness to genotoxic stress demonstrated in this study likely reflects their frontline activity in maintaining homeostasis. In contrast, the strong transcriptional activation observed in vibratile cells is more unexpected, as these cells are less abundant in whole coelomic fluid and are generally not well characterized. Pathway enrichment analysis of significantly upregulated genes in UVB-treated vibratile cells revealed many Reactome pathways related to cell cycle progression and mitosis. However, as the genes assigned to these pathways (such as *CCNE1, MCM8, GINS, DNA2,* and *POLD2*) are also involved in the replication stress response, we interpret this finding as the activation of DNA repair and replication stress response pathways, rather than active cell division. This is consistent with the limited proliferative capacity of circulating coelomocytes, which are considered to be largely post-mitotic (108). The vibratile cell response to UVB suggests a previously unrecognized role in response to genotoxic stress.

Red and colorless spherule cells exhibited only a minor transcriptomic response to UVB. Red spherule cells contain echinochrome A (EchA), a pigment that acts as an antioxidant (109–111). Consequently, the ability of EchA to quench reactive oxygen species generated by UVB may have conferred protection to these cells, resulting in the observed muted transcriptional response. In addition, *Bcl-2* genes were detected only in UVB-treated red spherule cells using scRNAseq, but were not detected in our bulk RNAseq analysis. This likely reflects the enhanced resolution of scRNAseq, which captures cell-type-specific signals that may be diluted in bulk RNAseq, in which gene expression is averaged across all cell types. The observed upregulation of *Bcl-2* genes only in red spherule cells suggests that apoptosis may be occurring only in this cell type, rather than broadly across all coelomocyte types, though additional validation using cell-type-specific assays is needed to validate this. Colorless spherule cells also exhibited a muted transcriptional response to UVB, although these cells do not express EchA and therefore lack any hypothetical pigment-based protection. Their low level of response may instead reflect a higher activation threshold or an intrinsic difference in sensitivity to DNA damage. Overall, these findings highlight the functional diversity of coelomocyte subtypes and their responses to genotoxic stress.

### 4.3 *TRIM, NLR, and RLR* gene expression is upregulated in response to UVB challenge

Among the immune genes found to be significantly upregulated in response to UVB-challenge, the majority of immune effectors belonged to the tripartite motif (TRIM) family. TRIM proteins are involved in various cellular processes, including cell differentiation, development, programmed cell death, and innate immunity (99). Most, but not all TRIM proteins function as E3 ubiquitin ligases mediated by a catalytic site within their RING domain. They function to polyubiquitinate target proteins, thereby regulating proteasome-mediated degradation, protein stability, and gene expression (via ubiquitination of transcription factors), among other functions (99,112,113). Within the context of innate immunity, TRIM proteins regulate PRRs and are also involved in antiviral responses via direct viral restriction, modulation of immune signaling, and autophagy of virally infected cells (114,115). TRIM proteins are present in all metazoans, although the number of family members varies among species. Interestingly, and in contrast with humans (68 family members), mice (67 family members), *Caenorhabditis elegans* (7 members), and *Drosophila melanogaster* (18 members) (116), the TRIM superfamily is expanded in *S. purpuratus*, with 140 genes annotated as “tripartite motif-containing protein” within Echinobase (117). Among the significantly expressed *TRIM* genes in UVB-challenged *S. purpuratus* phagocytes, *TRIM45*, *TRIM71*, *TRIM33*, *and TRIM56* were highly represented, with 10, 7, 5, and 2 unique upregulated genes, respectively (Supplementary Table 8). Notably, *TRIM* gene expression in vibratile cells was restricted to *TRIM45* genes (9 unique upregulated genes). Functional characterization of TRIM proteins has largely been conducted in vertebrate systems, specifically in human cell line and murine models, and several family members have been implicated in tumor suppression and innate immunity (112,118). For example, in humans, TRIM45 stabilizes and activates p53 in glioma models, functioning as a tumor suppressor (119), while TRIM33 (also known as TIF1γ) acts as a tumor suppressor in hepatocellular carcinoma, chronic myelomonocytic leukemia, and pancreatic cancer (120–123). TRIM56 has been linked to cytosolic DNA sensing and downstream NF-κB signaling via STING and TLR3 pathways (124,125), while TRIM71 is associated with stem cell proliferation and developmental regulation (126). Although functions of these *TRIM* gene orthologs have not yet been established in sea urchins, their significant upregulation following UVB challenge is consistent with the broader functions of TRIM proteins in cellular responses to DNA damage, immune activation, and maintenance of genomic integrity. Given the expansion of the *TRIM* gene family in sea urchins, this response may reflect an evolutionarily conserved strategy for responding to genomic stress or a lineage-specific adaptation that has diversified TRIM-associated functions. Future functional studies are expected to determine whether sea urchin TRIM proteins participate in signaling pathways similar to those described in vertebrates, or whether they have acquired novel functions in responding to DNA damage.

PRRs were significantly expressed in response to UVB-challenge, especially in phagocytes. NOD-like receptors (NLRs) were the most abundant among all PRR types. Sea urchins possess a rich repertoire of *NLR* genes within their genomes, with more than 200 *NLR* genes compared to only ∼20 in vertebrates (28,30). Many of these NLR genes contain NACHT and LRR domains, and in some cases, CARD-like domains. These features suggest functional parallels with vertebrate *NLR* genes, particularly those involved in inflammatory and stress responses. Overall, the upregulation of *NLR* genes in response to DNA damage indicates a role for NLRs in the DNA damage response in sea urchins. Enhanced expression of three *RIG-I* genes (also known as DExD-box helicase *DDX58* genes) was also observed in response to UVB. These genes are members of the RIG-I-like receptor (RLR) family. This family of RNA helicases is involved in the antiviral response, acting as viral nucleic acid sensors and regulating signal transduction downstream of various PRRs (127). RIG-I detects viral RNA in the cytoplasm and triggers type I interferon responses through the mitochondrial antiviral signaling protein MAVS (128,129). In addition to its well-characterized function as a sensor of viral infection, RIG-I also detects cytoplasmic mitochondrial RNA and triggers an immunological response (130), linking innate immune signaling to RNA sensing pathways. Sensors of mtRNA and cytosolic RNA are therefore crucial mediators between the immune system and DNA damage (13). Future work to characterize the proteins encoded by the *RIG-I* genes in sea urchins is expected to reveal their role in the DDR.

## 5 Conclusions

Our findings demonstrate that *S. purpuratus* coelomocytes respond to UVB-induced DNA damage with upregulated transcription of pattern recognition receptor genes, *TRIM* genes, and genes involved in DDR pathways and autophagy, suggesting an intricate coordination between DNA repair and innate immunity. Due to their remarkable longevity, absence of reported cases of neoplasia, accessible and manipulable immune system, and lack of canonical vertebrate DNA sensing pathways such as cGAS-STING, sea urchins represent a unique model system that can be leveraged to uncover novel mechanisms for preserving genomic integrity. These findings lay the foundation for further studies into evolutionarily conserved and lineage-specific DDR-immune interactions in early deuterostomes and their function in shaping organismal resilience and protection from neoplasia.

## Data availability statement

Raw data for the single-cell and bulk RNA sequencing have been deposited in the National Center of Biotechnology Information (NCBI) under BioProject accession number PRJNA1346447 (Sequence Read Archive accession numbers SRR35853817 – SRR35853834). Code used for transcriptomic data analysis and visualization are available at: https://github.com/rmkell/Spurpuratus-immunesystem-DNAdamage/. Other data presented in this study are included in the Supplementary Materials.

## Author contributions

RMK: Conceptualization, Data curation, Formal analysis, Investigation, Methodology, Visualization, Writing – original draft, Writing – review & editing. MW: Investigation, Formal analysis, Visualization, Writing – review & editing. RE: Investigation, Formal analysis, Visualization, Writing – review & editing. JS: Conceptualization, Methodology, Resources. AGB: Funding acquisition, Conceptualization, Resources, Supervision, Project administration, Writing – review & editing.

## Acknowledgements

We thank Dr. Kate Castellano and Jennifer Polinski for assistance with scRNA-seq training and in validating cDNA libraries, Dr. Bo Reese for assistance with Illumina sequencing, Dr. Courtney Smith for providing feedback on the manuscript, Dr. Emma Strand for guidance regarding bulk RNAseq analysis, and Reanna McAtee for assistance with sea urchin maintenance and coelomocyte sampling.

