## Supplementary Figures for "UVB-Induced Genotoxic Stress Activates the DNA Damage Response and Innate Immune Pathways in Sea Urchin Coelomocytes"

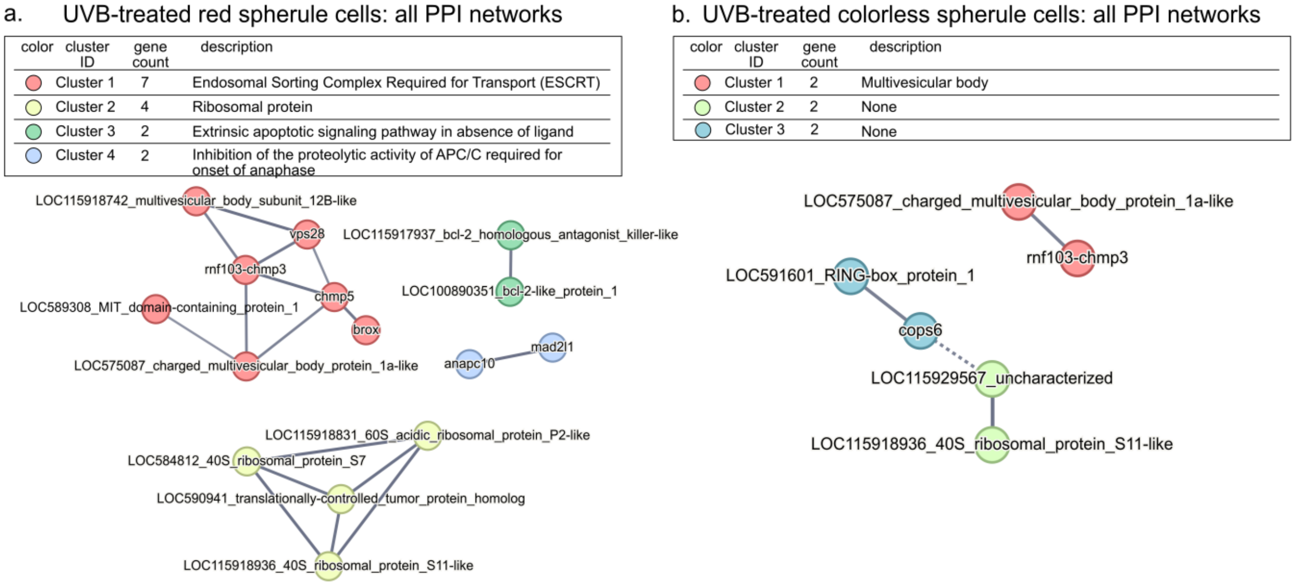


**Supplementary Figure 1. Only minimal protein-protein-interaction (PPI) networks were detected among proteins encoded by significantly upregulated genes in UVB-treated red and colorless spherule cells.** Graphical representation of all protein-protein interactions (determined by k-means clustering) among proteins encoded by the significantly upregulated genes ($p < 0.05$, $\log_2$foldchange > 1) using the STRING database (v12.0) with high-confidence settings. (a) PPI networks among proteins encoded by upregulated genes in UVB-treated red spherule cells. (b) PPI networks among proteins encoded by significantly upregulated genes in UVB-treated colorless spherule cells. Complete lists of enriched terms are given in Supplementary Table 7.

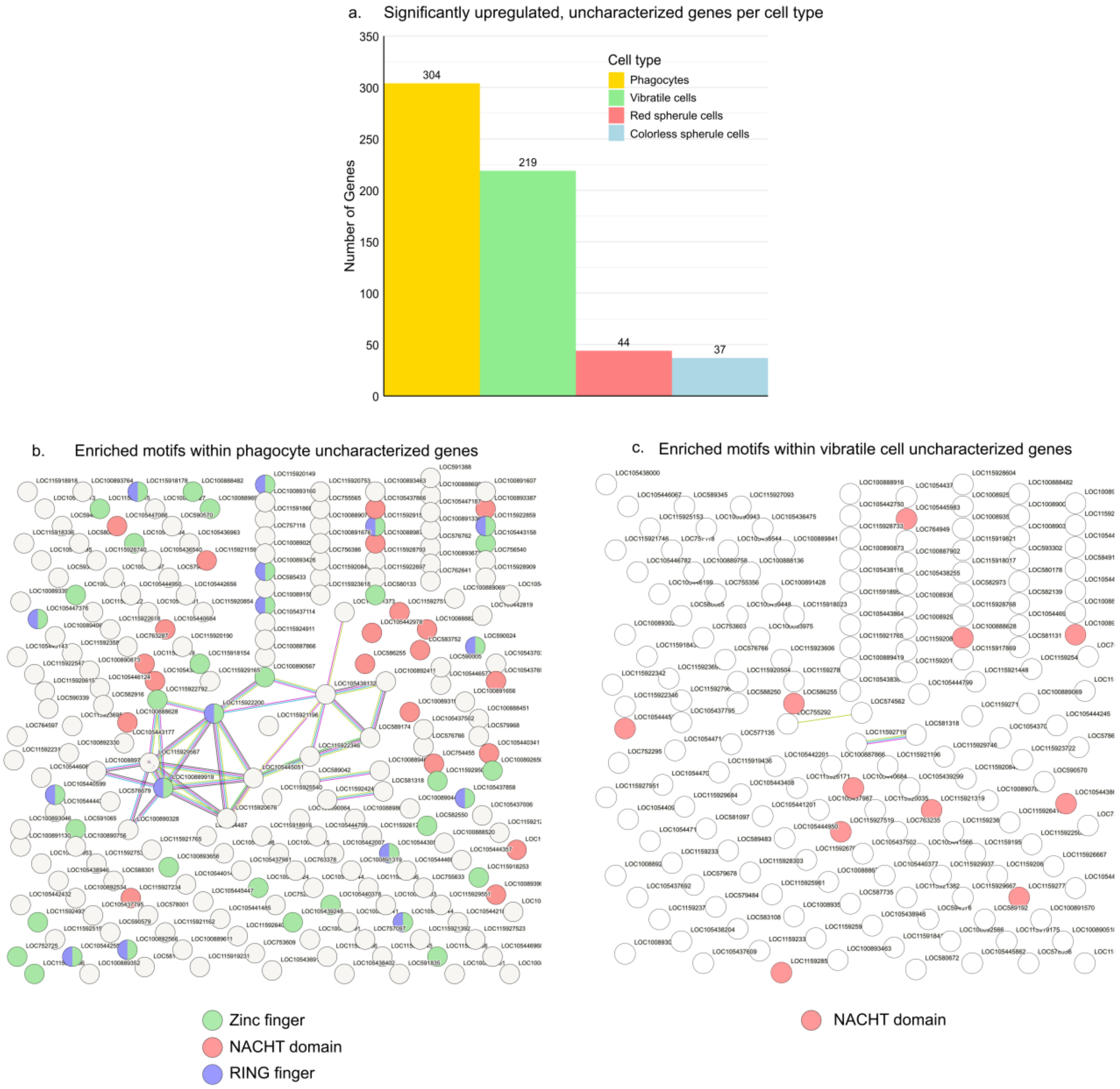


**Supplementary Figure 2**. **Upregulated genes encoding proteins annotated as "uncharacterized" are enriched in zinc finger motifs, NACHT domains, and RING finger motifs.** (a) Bar chart quantifying significantly upregulated ($p < 0.05$, $\log_2$foldchange > 1) genes annotated as "uncharacterized" in UVB-treated cells compared to control cells by cell type. (b) STRING analysis of the proteins encoded by the 304 uncharacterized genes in UVB-treated phagocytes shows enrichment of zinc finger motifs, NACHT domains, and RING finger motifs. (c) STRING analysis of the proteins encoded by the 219 uncharacterized genes in UVB-treated vibratile cells shows enrichment of NACHT domains. No significant enrichment was detected among the proteins encoded by the significantly upregulated, uncharacterized genes in red or colorless spherule cells. STRING networks were generated under medium confidence settings. A complete list of enriched terms is given in Supplementary Table 9.

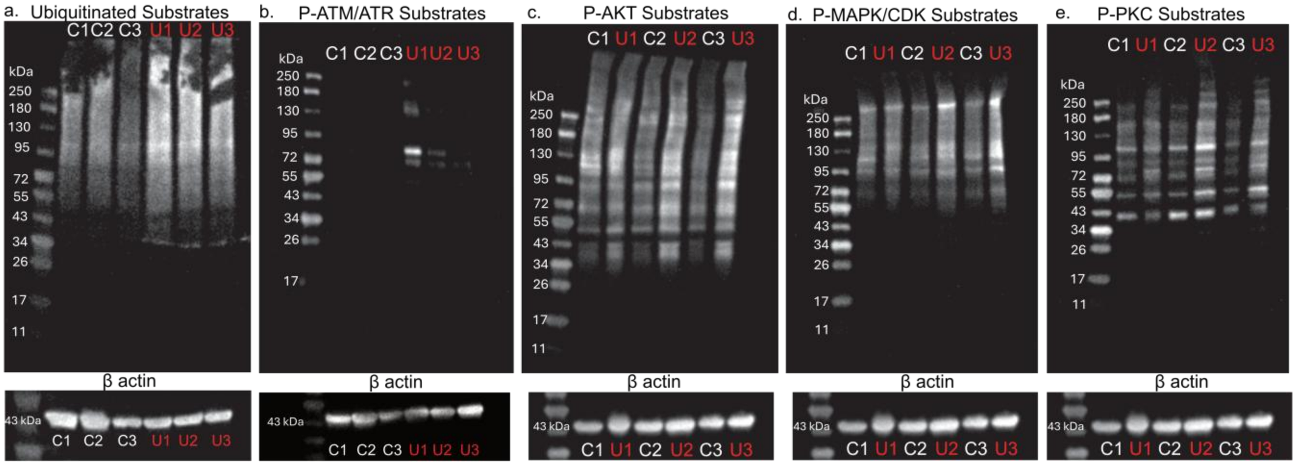


**Supplementary Figure 3. Additional western blots.** Western blots showing (a) ubiquitinated, (b) phosphorylated ATM/ATR substrate, (c) phosphorylated AKT substrate (d), phosphorylated MAPK/CDK substrate, and (e) phosphorylated PKC substrate signal in biological triplicate control (C1-C3, white font) and UVB-treated (U1-U3, red font) samples. β actin loading controls are shown below. Quantification of western blot signals is presented in Figure 9 in the main paper. The dark blotches in the ubiquitin blot (a) was the outcome of the SDS-PAGE gel that melted and adhered to the nitrocellulose membrane during transfer. Signal quantification did not include the blotches; the rectangular ROIs began just under 180 kDa). The ubiquitin blot presented in Figure 9a in the main paper was quantified in the same way for consistency. Blots run simultaneously (the P-AKT, P-MAPK/CDK, and P-PKC blots presented in this figure, and the P-ATM/ATR blot shown here and the P-AKT blot shown in the main text in Figure 9e) have identical β actin loading controls.

### Supplementary Table legends

**Supplementary Table 1**. **Differentially expressed genes revealed by timecourse bulk RNA sequencing.** Tab 1: BulkRNAseq: differentially expressed (Benjamini-Hochberg adjusted *p* values < 0.05, $\log_2$ fold-change < -1 or > 1) genes comparing control and UVB-treated samples at each timepoint (0, 1, 3, 6, 24 h). Each gene was assigned to a k-means cluster (Cluster 1 or Cluster 2).
Tab 2: All significantly enriched ($p < 0.05$) STRING categories (GO, KEGG, Reactome, Compartments, UniProtKeywords, SMART domains) assigned to PPI networks found among Cluster 1 genes, generated using highest confidence STRING settings.
Tab 3: All significantly enriched ($p < 0.05$) STRING categories (GO, KEGG, Reactome, Compartments, UniProtKeywords, SMART domains) assigned to PPI networks found among Cluster 2 genes, generated using highest confidence STRING settings.
Tab 4: Gene products upregulated in UVB-treated samples at all timepoints.
Tab 5: Gene products downregulated in UVB-treated samples at all timepoints.
Tab 6: Summary of the number of up- and down-regulated genes in UVB-treated samples over time.

**Supplementary Table 2**. **ScRNAseq total reads**. Total cells and fraction of reads in cells in the raw scRNAseq matrix HD files before and after preprocessing.

**Supplementary Table 3. Fraction of control or UVB-treated cells in each Leiden cluster.** Data on the first tab, "PercentConditionbyCluster", lists the Leiden cluster number, the percentage of cells in each condition (control or UVB-treated) comprising that cluster, and the cell type assigned to that cluster. Green highlighting indicates those clusters that are predominantly (> 80%) comprised of either control or UVB-treated cells. Yellow highlighting indicates those clusters that are comprised of a mix of both control and UVB-treated cells. Data on the second tab, 'Fraction Each CellType', shows the cell type fractions in each sample (data plotted in Fig. 3d).

**Supplementary Table 4. Differential expression of Leiden clusters and cell type assignment.** Assignment of scRNAseq leiden clusters to broad coelomocyte cell type categories (phagocytes, vibratile cells, colorless spherule cells, red spherule cells) based on marker genes from the literature. Genes were ranked based on their differential expression across leiden clusters using the Scanpy library 'sc.tl.rank_genes_groups' function. The ranked genes and their statistics were retrieved as a dataframe named 'markers'. The markers dataframe was filtered to keep only those genes with adjusted *p* values < 0.05 and $\log_2$ fold changes > 0.5. LOC gene IDs and descriptive names ('gene name' column) are provided. 'Scores', 'logfoldchanges', 'pvalue', and 'pval adjusted' columns refer to statistics generated from differential expression analysis of individual genes comparing expression among all clusters. Adjusted *p* values were corrected using the Benjamini-Hochberg procedure. Scores are the z-score underlying the computation of a *p* value for each gene in each group. Logfoldchanges are an approximation calculated from mean-log values.

**Supplementary Table 5. Gene Ontology (GO) categories of differentially expressed genes comparing all control cells to all UVB-treated cells.** Enriched Gene Ontology (GO) terms assigned to significantly upregulated (Tab 2) and downregulated (Tab 3) genes in UVB-treated samples using GOATOOLS. GO: the Gene Ontology numerical identifier. Term: GO term description. N_genes: The number of genes associated with the GO term in the entire dataset (plotted in Fig. 2). Class: GO term category. P: p-value of GO term enrichment. P_corr: Corrected *p* value after false discovery rate (FDR) correction using the Benjamini-Hochberg procedure. N_study: the total number of genes in the study dataset. N_go: the total number of genes in the GO term. The background gene set used with GOATOOLS analysis was all NCBI *S. purpuratus* (NCBI taxonomy ID 7668) protein coding genes.

**Supplementary Table 6. Differentially expressed genes in UVB-treated versus control coelomocytes at the cell-type level.** Data used to generate the volcano plots shown in Fig. 5. Differential gene expression results comparing control and UVB-treated phagocytes, vibratile cells, red spherule cells, and colorless spherule cells are provided.

**Supplementary Table 7. STRING enrichment by cell type.** All significantly enriched ($p < 0.05$) STRING categories (GO, KEGG, Reactome, Compartments, UniProtKeywords, SMART domains) assigned to protein-protein-interaction (PPI) networks found among proteins encoded by significantly upregulated ($p < 0.05$, logfoldchange > 1) genes in UVB-treated cells compared to control cells, by individual cell type (phagocytes, vibratile cells, red spherule cells, and colorless spherule cells). STRING was run under high confidence settings. Tabs describe those k-means clusters with > 10 nodes. Column names are as follows: 'observed gene count', the number of proteins in the network that are annotated with a particular term; 'background gene count', the total number of proteins (in network plus in the background) annotated with a particular term; 'strength', log10(observed/expected), a measure that describes how large the enrichment effect is. It is the ratio between the number of proteins in the PPI network that are annotated with a term and the number of proteins expected to be annotated with this term in a random network of the same size. 'Signal': a weighted harmonic mean between the observed/expected ratio and -log(FDR). 'False discovery rate': describes how significant the enrichment is. Shown are p-values corrected for multiple testing within each category using the Benjamini–Hochberg procedure.

**Supplementary Table 8**. **Enriched immune genes by cell type**. Significantly upregulated ($p < 0.05$, $\log_2$foldchange > 1) immune system genes in UVB-treated cells compared to control cells, by individual cell type (phagocytes, vibratile cells, red spherule cells, colorless spherule cells), assigned to immune gene categories.

**Supplementary Table 9**. STRING analysis of significantly upregulated ($p < 0.05$, $\log_2$foldchange > 1) uncharacterized genes in UVB-treated cells compared to control cells, by individual cell type, showing significantly enriched ($p < 0.05$) terms.
